# Microbial Priming Enhances TLR7-Driven Myddosome Assembly Dynamics

**DOI:** 10.64898/2025.12.16.693659

**Authors:** Anna Dingfelder, Fenja Blank, Noemie Quinson, Caroline Holley, Jonathan Mayr, Oliver Thieck, David Meierhofer, Marcus J Taylor, Olivia Majer

## Abstract

Cells can modify their future behavior based on prior exposures, a phenomenon known as cellular adaptation. In innate immunity, such adaptive responses are critical for fine-tuning host defense, enabling trained immunity or tolerance. These states have been largely attributed to long-lasting epigenetic or metabolic reprogramming. Whether prior microbial encounters can rapidly lower receptor signaling thresholds to selectively enhance responsiveness of individual innate immune receptors remains unknown.

Here, we identify a previously unrecognized, non-genetic form of cellular adaptation. We show that TLR4 activation by LPS, while inducing classical endotoxin tolerance, simultaneously primes macrophages for enhanced responses to subsequent activation of the RNA sensor TLR7. Because TLR4 and TLR7 use the same signaling machinery, this indicates that they are selectively reprogrammed in this process. TLR7 training requires type I interferons and is marked by a strong increase in Myddosome assemblies at endosomal membranes, without changes in TLR7 abundance or ligand uptake.

These findings reveal that innate immune training can be rapidly encoded at the level of receptor-proximal signaling, linking microbial priming to enhanced nucleic acid immunity and potentially to TLR7-driven autoimmunity in genetically predisposed individuals.

**Short summary:** LPS priming drives receptor-proximal immune adaptation in macrophages, inducing classical TLR4 endotoxin tolerance while simultaneously sensitizing the RNA sensor TLR7 by enhancing Myddosome nucleation at endosomal membranes. This compartment- and receptor specific reprogramming may link prior infection history to elevated risk of TLR7-driven autoimmunity.

## Introduction

In response to environmental cues, cells can modify their properties in ways that influence future responses to the same or other stimuli, a process known as cellular adaptation (Davies, 2016). This phenomenon enables cells to adjust their functional state to match environmental demands, maintaining core processes while optimizing responses to recurring or novel stimuli. While adaptation represents a widespread phenomenon across diverse physiological processes, it remains unclear how prior exposures can rapidly reconfigure signaling networks to alter the responsiveness of individual receptors.

The innate immune system provides a prime physiological context in which cellular adaptation is used to scale and tailor responses for effective host defense. During repeated or sequential infectious encounters, immune cells are exposed to dynamic and overlapping signals. These signals must be properly integrated, so that subsequent challenges are either tolerated or cleared with a heightened response. Such innate immune adaptations are the basis for enhanced protection against unrelated pathogens after a prior infection or vaccination, a phenomenon termed innate immune memory or *trained immunity* (Netea et al., 2020). In contrast, prior exposure to certain ligands, such as bacterial lipopolysaccharide (LPS), can induce *tolerance*, characterized by reduced cytokine output (West and Heagy, 2002). These adaptive immune states have been primarily attributed to long-term epigenetic and metabolic changes. However, it remains unknown whether and how cellular adaptation is also encoded at the level of receptor-proximal signaling networks to induce a rapid and stronger response after a primary stimulus.

Cellular adaptation may be particularly critical when increased immune responses can have detrimental effects for the host, as in the case of nucleic-acid sensing Toll-like receptors (TLRs). These endosomal receptors detect microbial nucleic acids and play central roles in antiviral defense. However, tight regulation of their activity is essential to prevent inappropriate responses to self-derived nucleic acids, and failure of this regulation can lead to autoimmunity. This is particularly true for the RNA sensor TLR7, for which hyperactive responses have long been appreciated as a driver of autoimmune disease in mouse models (Deane et al., 2007; Fukui et al., 2011; Majer et al., 2019), and more recently in humans (Brown et al., 2022; Mishra et al., 2024; Rael et al., 2024; Wolf et al., 2024).

While the genetic basis of TLR7-driven autoimmunity is increasingly appreciated, the interplay between environmental, infectious and genetic factors common to autoimmune diseases suggests an additional role for cellular adaptation. Notably, autoimmune flares in susceptible individuals frequently correlate with prior infections, implicating a potential priming effect by microbial products (Battaglia and Garrett-Sinha, 2021; Doria et al., 2008). In patients with a predisposition for TLR7-driven autoimmunity, such priming could transiently lower the threshold for TLR7 signaling, promoting the recognition of self-derived nucleic acids. However, how pathogenic cues might sensitize endosomal TLRs to nucleic acids, and how this adaptation occurs at a molecular level, remain unknown.

In this study, we describe a previously unrecognized form of cellular adaptation in macrophages, in which microbial ligands reprogram receptor-proximal signaling dynamics to sensitize TLRs to nucleic acid ligands *in vitro*. Specifically, we show that TLR4 activation, while inducing classical endotoxin tolerance, simultaneously primes macrophages for enhanced responses to TLR7 stimulation. This training requires type-I interferons and is marked by a strong increase of MyD88 signalosome assemblies at endosomal membranes. As a result, receptor activation leads to enhanced MAP kinase phosphorylation and cytokine production. These findings reveal that prior microbial exposure can differentially modulate the kinetics of shared signaling pathways downstream of TLRs, enabling faster and stronger inflammatory responses. It suggests that, in the case of TLR4 and TLR7, training and tolerance may occur simultaneously in a receptor-specific manner. Our results expand the concept of trained immunity beyond transcriptional and metabolic reprogramming, uncovering a cell-intrinsic, non-genetic mechanism that encodes innate immune adaptation at the level of signal transduction. Moreover, this mechanism provides a plausible molecular framework linking infection-induced immune priming to the breach of tolerance toward self-derived nucleic acids, offering new insights into the potential origins of TLR7-driven autoimmunity in genetically predisposed individuals.

## Results & Discussion

### 1. LPS priming amplifies signaling responses of nucleic acid-sensing TLRs

To understand how cellular adaptation is encoded at the molecular level within the innate immune system, we assessed the priming effect of pathogenic cues on nucleic acid-sensing Toll-like receptors. The TLR family shares the same molecular signaling machinery across different compartments, allowing for direct comparison of their kinetics after a priming stimulation. We focused on macrophages, given their role as tissue sentinels that sense both environmental and immunological signals (Davies and Taylor, 2015; Wynn et al., 2013). They are known to adapt their function to diverse conditions and tissue contexts, often supporting long lasting adaptations (Stout and Suttles, 2004). Macrophages therefore present a suitable model system to investigate the mechanisms of cellular adaptation rooted in dynamics of immune signaling pathways.

Based on previous studies on trained immunity, we adopted a stimulation regimen in which a primary stimulation is followed by a secondary challenge with nucleic acid ligands. We first primed macrophages derived from Hoxb8-immortalized bone marrow progenitors (referred to as ‘Hoxb8 macrophages’; (Wang et al., 2006) for 24h with a variety of TLR ligands, including LPS (TLR4), Pam3CSK4 (TLR2), R848 (TLR7) and CpG-B (TLR9). Several of these ligands are widely used in the trained immunity field and they are thought to mimic viral or bacterial infection. We made sure that 1) each priming stimulus activated the entire macrophage population, by reading out TNF production at the single-cell level 6 hours post priming, and 2) TNF response returned back to baseline after 24 hours. The priming stimulus was then rigorously washed out, and cells were re-stimulated with the second ligand (Fig. 1A). This system allowed to screen different combinations of primary and secondary stimulation within single cells, identifying ligand interactions resulting in a trained or tolerant immune response, as measured by intracellular TNF staining (Fig.1B). We found that, in each case, a ligand induced tolerance to itself upon re-challenge. This behavior is best described for TLR4/TLR4, also known as endotoxin tolerance (Seeley and Ghosh, 2017). Moreover, we were able to reproduce previously reported cross-tolerance between other ligand combinations, such as TLR2/TLR4, and TLR2/TLR9 (Dalpke et al., 2005). These findings demonstrate the validity of our system to interrogate the molecular mechanisms underpinning short-term cellular adaptations.

**Figure 1.**
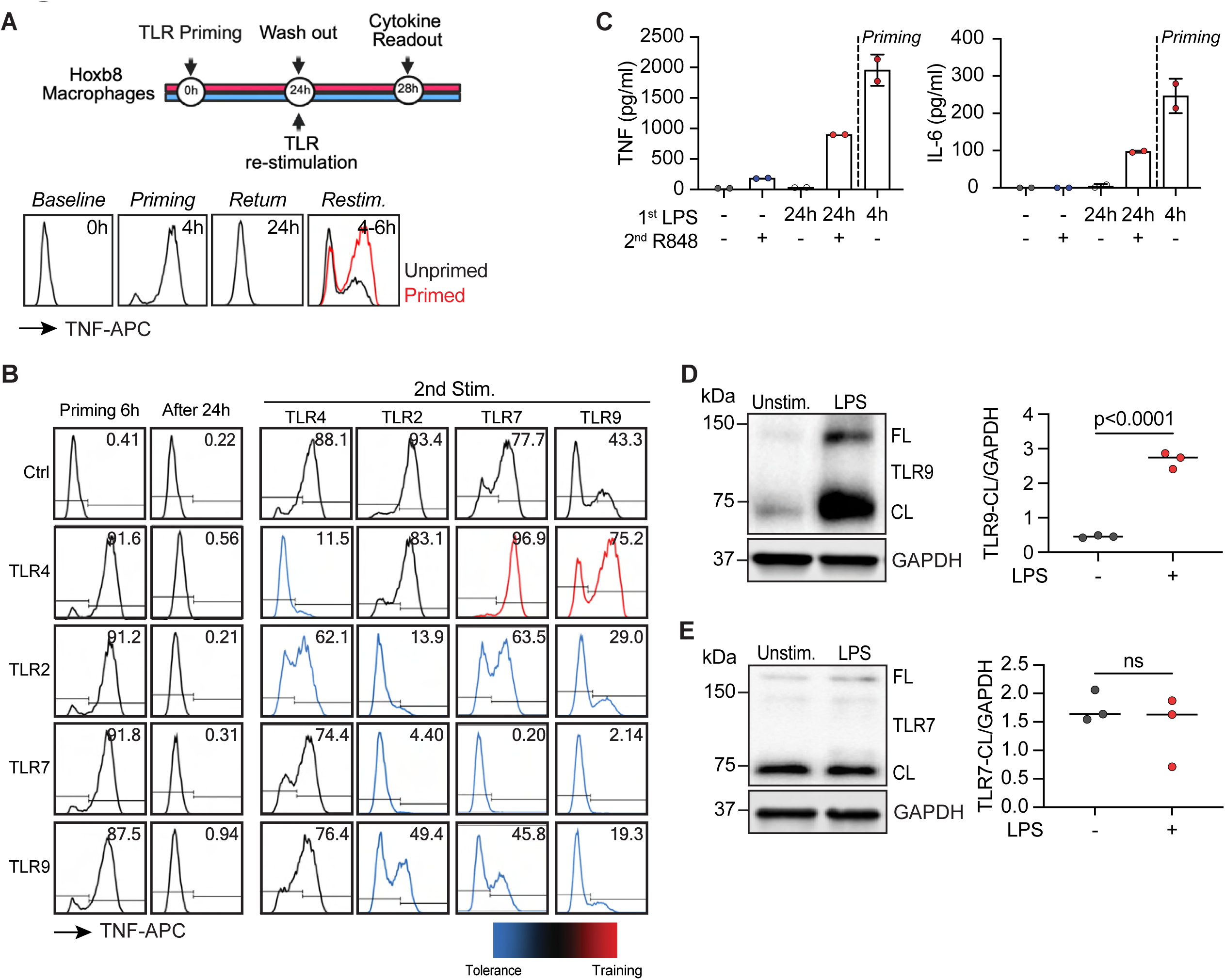
LPS priming enhances TLR7 signaling without receptor upregulation. **(A)** Priming schematic for Hoxb8 macrophages. (**B)** Intracellular TNF staining of primary/secondary TLR stimulations. Cells were primed with TLR ligands: LPS 100ng/ml (TLR4), Pam3CSK4 200ng/ml (TLR2), R848 100ng/ml (TLR7) and CpG-B 100nM (TLR9). Priming ligand was washed out after 24h and cells were re-stimulated with secondary TLR agonists: LPS 100ng/ml (TLR4), Pam3CSK 100ng/ml (TLR2), R848 10ng/ml (TLR7) and CpG-B 50nM (TLR9). Colors indicate observed effect: red = enhanced response upon secondary stimulation (training), blue = reduced response (tolerance). Black = unchanged. Representative experiment out of N=3. (**C)** Legendplex assay of TNF and IL-6 (N=2) of cells stimulated as in (B). **(D)** Western blot analysis of TLR7 protein levels after LPS priming (N=3). **E:** Western blot analysis of TLR9 protein levels after LPS priming (N=3). CL: cleaved. P values were calculated with ratio paired t-test. ns: not significant.

After confirming these known effects, we explored potential training constellations of cell surface and endosomal nucleic-acid sensing TLRs. Strikingly, we found a unique and previously unrecognized effect of LPS exposure on subsequent responses to nucleic acids. While LPS-priming had a tolerizing effect on subsequent LPS stimulation, it simultaneously trained TLR7 and 9 responses, as evidenced by increased TNF and IL-6 production upon secondary challenge (Fig. 1B, C). This suggests that prior bacterial exposure sensitizes subsequent nucleic acid sensing within a short time frame. Enhanced TLR7 responses after LPS priming were also observed in murine *ex vivo* alveolar macrophages (Supp. Fig.1A). We therefore conclude that TLR4 stimulation induces a unique cellular adaptation that selectively trains TLR7 and TLR9 - receptors that share the same proximal signaling machinery yet respond oppositely to prior pathway activation.

To assess whether this enhanced response was due to transcriptional upregulation of TLR7/9 after priming, we performed western blot analysis. While LPS priming strongly upregulated TLR9 expression, TLR7 levels remained unaffected (Fig. 1D, E, Supp. Fig. 1B, C). This indicates that LPS priming differentially regulates the expression of endosomal TLRs and that increased TLR7 signaling is not a consequence of increased receptor abundance but rather enhanced activation. To rule out increased ligand uptake after LPS priming, we compared the internalization of fluorescent CpG-B-Cy5 between naive and LPS-primed macrophages for up to 3h. Flow cytometry revealed no difference in ligand uptake in primed cells (Supp. Fig. 1D).

Because TLR7 gain-of-function is strongly associated with loss of self-tolerance and autoimmunity (Brown et al., 2022), we decided to explore this training effect further. Titration of the primary and secondary stimulation revealed that trained TLR7 responses were most apparent at low R848 concentrations (Supp. Fig.1E). Training of TLR7 was most effective at an LPS priming concentration of 100ng/ml and declined with decreasing concentrations (Supp. Fig.1F). Induction of endotoxin tolerance was also most effective at 100 ng/ml LPS, with tolerance decreasing as the priming dose was reduced (Supp. Fig.1G).

In summary, we show that TLR4 stimulation not only induces the well-known state of endotoxin tolerance but also concurrently sensitizes the nucleic acid sensors TLR7 and TLR9, with both of these opposing outcomes converging on the same proinflammatory cytokines. This indicates that LPS-tolerized cytokine genes such as *Tnf* and *Il6* are not subject to uniform global suppression during prolonged LPS exposure, but are instead differentially controlled by the upstream receptor and intracellular signaling context. These findings challenge the prevailing view that tolerance and training are mutually exclusive, global cellular states (Divangahi et al., 2021; Lajqi et al., 2023). Rather, our data support a model in which a single priming stimulus elicits opposing functional adaptations in distinct receptor compartments that use the same signaling machinery, attenuating signal transduction and gene expression from the cell surface while enhancing it from endosomes in a receptor-specific manner.

### 2. TLR7 training depends on TLR4-TRIF-IRF3 signaling and type I interferons

To determine which signals drive enhanced activation of TLR7 after priming, we systematically delineated the signaling events downstream of TLR4 activation. Upon TLR4 engagement, two pathways are activated: the MyD88-dependent pathway inducing proinflammatory cytokines and the TRIF-IRF3-dependent pathway that drives type I interferon (IFN-I) production (Supp. Fig. 2A).

IFN-I has previously been shown to play a critical role in LPS-induced trained immunity in alveolar macrophages, conferring enhanced protection against pneumococcal challenge *in vivo* (Zahalka et al., 2022). To test whether IFN-I was involved in TLR7 training, we used CRISPR-Cas9 genome editing to knock out components of the TRIF-dependent signaling pathway that either mediate IFN-I production (TRIF, IRF3) or its autocrine/paracrine sensing (IFNAR2) (Supp. Fig. 2A). Knockouts were functionally validated by measuring production of the interferon-stimulated gene (ISG) Viperin upon LPS stimulation via intracellular flow cytometry (Supp. Fig. 2B). As expected, knockout of TRIF or the transcription factor IRF3 resulted in loss of Viperin production, and knockout of the interferon receptor subunit IFNAR2 also abrogated Viperin expression. Viperin production could be rescued in TRIF^-/-^ and IRF3^-/-^ (but not IFNAR2^-/-^) macrophages by stimulating with recombinant murine IFN-β (Supp. Fig. 2B). We further confirmed that these knockout lines responded to the initial LPS priming stimulation similarly to wild type cells by measuring TNF production (Supp. Fig. 2C).

Strikingly, the increased response of TLR7 after LPS priming was completely abrogated in all three knockout lines after LPS priming, as evidenced by reduced IL-6 induction upon R848 restimulation (Fig. 2A-C). This data indicate that IFN-I must be both secreted and sensed during LPS priming to confer heightened TLR7 sensitivity. Priming could be rescued in IRF3^-/-^macrophages by supplementing recombinant murine IFN-β during the LPS priming phase (Fig. 2D). However, IFN-β priming alone was not sufficient to induce training (Fig. 2E). Likewise, supernatant transfer experiments, in which residual LPS from primed cells was neutralized with Polymyxin-B, failed to confer training to naive recipient cells (Fig. 2F, Supp. Fig. 2E). Collectively, these results demonstrate that soluble mediators alone are not sufficient to induce TLR7 training and that additional cell-autonomous signals triggered by LPS are required. This is consistent with cellular adaptation requiring both IFN-I signaling and additional TLR4-dependent signals acting in concert to confer TLR7 training.

**Figure 2.**
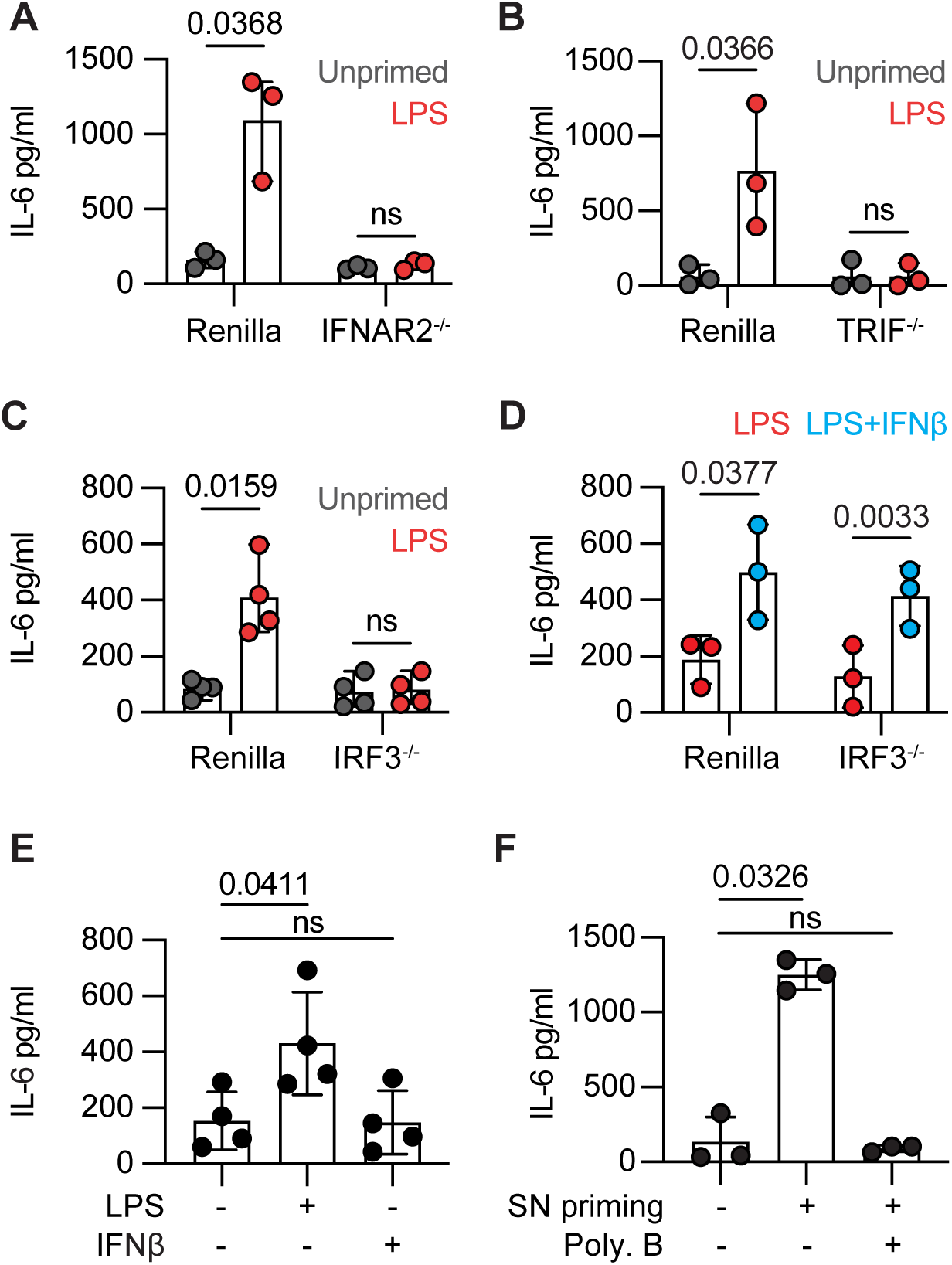
TLR7 training depends on TLR4-TRIF-IRF3 signaling and IFNs-I. **(A-C)** IL-6 ELISA of IFNAR^-/-^, TRIF^-/-^ and IRF3^-/-^ macrophages. Cells were primed with LPS 100ng/ml and restimulated with R848 10ng/ml (N=3). **(D)** IL-6 ELISA of WT and IRF3^-/-^ macrophages primed with LPS 100ng/ml and IFN-b 800U/ml (N=3). **(E)** IL-6 ELISA of WT macrophages primed with IFN-b 800U/ml (N=3). **(F)** IL-6 ELISA of supernatant (SN) transfer: WT Hoxb8 macrophages were primed with LPS 100ng/ml for 24h; supernatant was then transferred to naive cells to prime subsequent TLR7 activation. Neutralization of residual LPS in the supernatant with 50µg/ml Polymyxin B prevented priming of naive cells. Graph shows pooled independent repeats (N=3). (A-D): P-values were calculated with paired t-test. (E-F): P-values were calculated with one-way ANOVA with Tukey’s post-test. ns: not significant.

To understand the TLR4-dependent signaling required for TLR7 training, we sought to rule out the that LPS priming required the MyD88-NFκB signaling axis. We inhibited Myddosome assembly with the IRAK4 inhibitor Zimlovisertib, which prevents proinflammatory cytokine production downstream of surface TLR4 and endosomal TLRs (Supp. Fig. 2D) (De Nardo et al., 2018). To assess whether IRAK4 activation is required during the priming phase, we stimulated macrophages with LPS in the presence of 20µM inhibitor. After 24h, cells were rigorously washed, rested for 2h, and re-stimulated with R848 in the absence of the inhibitor, ensuring that residual inhibitor would not interfere with TLR7 signaling. As measured by IL-6 ELISA, TLR7 training was not reduced when Zimlovisertib was present during TLR4 priming (Supp. Fig 2F), even though the acute LPS priming response was markedly reduced at this inhibitor concentration (Supp. Fig. 2D). Thus, TLR7 training is conferred through a cell-autonomous TRIF-IRF3 stimulation by TLR4.

To test whether other IRF3-engaging receptors could mediate TLR7 priming, we stimulated macrophages with Poly:IC, a TRIF-dependent TLR3 agonist. Similar to LPS, Poly:IC priming sensitized macrophages to R848 without upregulating TLR7 expression, reinforcing the notion that TRIF-IRF3 signaling and IFN-I together are essential for this priming effect (Supp. Fig. 2G-H). Therefore, we conclude that TRIF-IRF3 are the essential signaling machinery by which TLR4 induces the cellular adaptation of TLR7 training.

The requirement for IFN-I therefore provides a molecular link between microbial exposure and increased endosomal TLR sensitivity. Together with cell-autonomous signals, IFN-I acts as a critical intermediary that induces a unique state of cellular adaption in which TLR7 is sensitized while TLR4 is desensitized, despite both receptors signaling through the same downstream machinery. Our findings are consistent with TLR7 training occurring at the level of receptor-proximal signaling.

### 3. LPS priming enhances MAPK phosphorylation downstream of TLR7 and 9

To distinguish whether the enhanced signaling output after LPS priming arises from epigenetic changes on cytokine transcription or from changes in the signaling threshold of TLR7, we assessed the immediate signaling response downstream of TLR activation. MAP kinases mark a central pathway in transducing TLR signals into pro-inflammatory gene expression. Using phospho-specific antibodies, we measured the kinetics and amplitude of MAPK activation following TLR7, TLR9 and TLR4 stimulation in either naive or LPS-primed macrophages.

LPS-primed cells exhibited stronger and more sustained phosphorylation of ERK1/2 and p38 MAPKs in response to TLR7 stimulation compared to unprimed cells (Fig. 3A). MAPK activation was also enhanced upon re-stimulation of TLR9 in primed cells with CpG-B, an effect likely attributable to its transcriptional upregulation (Fig. 3B). By contrast, MAPK activation was completely abolished upon restimulation of TLR4 (Fig. 3C), consistent with earlier studies reporting attenuated TLR4 signaling, in part due to the upregulation of negative-feedback regulators, during endotoxin tolerance (Kobayashi et al., 2002; Medvedev et al., 2000; Xiong et al., 2011). These findings indicate that LPS priming does not uniformly affect all TLR responses but instead selectively enhances signal transduction downstream of TLR7. Notably, this potentiation occurs early in the signaling pathway, upstream of cytokine transcription, suggesting a receptor-proximal mechanism.

**Figure 3.**
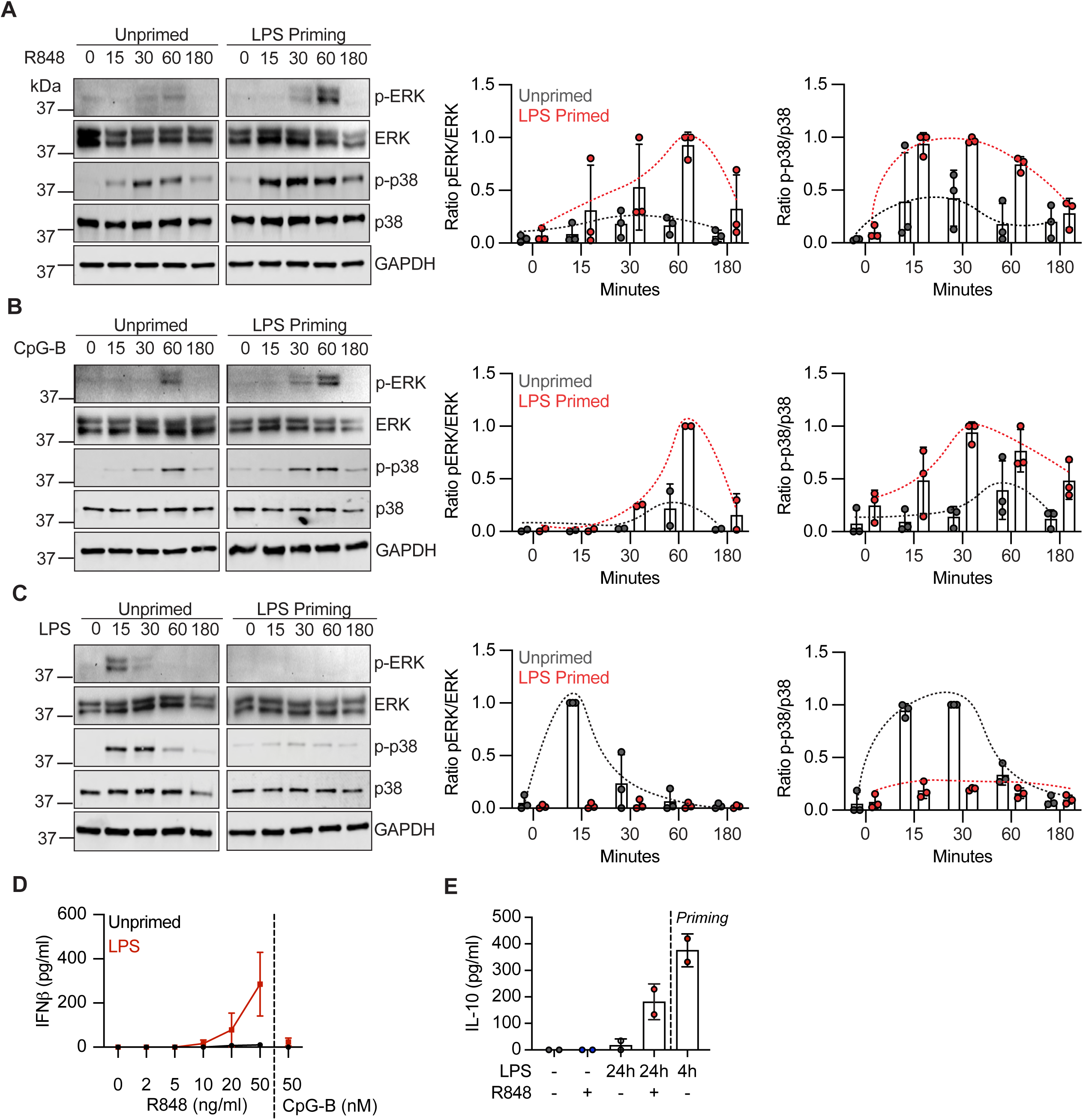
LPS priming enhances TLR7-dependent MAP kinase phosphorylation. **(A-C)** MAP kinase immunoblots of ERK and p38 phosphorylation upon TLR7, TLR9 or TLR4 stimulation in LPS-primed or naive cells. Representative blots out of N=3. Quantification performed on 3 pooled repeats. **(D)** Bioassay measuring IFN-b production by naïve or primed Hoxb8 macrophages upon TLR7/9 restimulation with increasing concentrations (N=3, shown is SEM). **(E)** Legendplex assay of IL-10 secretion: Hoxb8 macrophages were primed with LPS and re-stimulated with R848 10ng/ml. (N=2).

In human monocyte-derived dendritic cells, TLR4 stimulation has been shown to synergize with TLR7/9 to boost production of selected cytokines, including TNF, IFN-β, IL-12, IL-23, and IL-10, thereby promoting T helper type I responses (Napolitani et al., 2005). Analogous to this finding, LPS priming not only enhanced but also diversified the cytokine profile of TLR7-stimulated mouse macrophages. After 24h of LPS priming, R848 stimulation resulted in production of IFN-β and IL-10 (Fig. 3D, E), cytokines not expressed after TLR7 stimulation alone and typically associated with dendritic cells upon stimulation of endosomal TLRs. This suggests that receptor-specific signaling pathways are selectively re-wired after priming, or that heightened signaling output allows for transcriptional induction of otherwise poorly accessible cytokine genes. Collectively, these results indicate that LPS priming reconfigures the signaling landscape of macrophages, enabling faster and stronger immune activation from endosomal TLRs as well as expanding their functional plasticity and cytokine repertoire. We therefore conclude that LPS priming establishes a receptor-proximal adaptation that serves to amplify signal transduction through TLR7.

### 4. The MyD88-mNeonGreen knock-in mouse allows visualization of immune signaling

If TLR7 training is encoded within the receptor-proximal signaling network, it should manifest as altered assembly of functional signaling complexes upon ligand stimulation. To test this, we directly assessed Myddosome formation, which is the most proximate signaling event downstream of TLR activation. Activated TLRs recruit the signaling adaptor MyD88, which assembles with IRAK proteins into a multimeric signalosome termed the Myddosome (Deliz-Aguirre et al., 2021; Latty et al., 2018; Lin et al., 2010). To track this process, we employed a novel knock-in mouse model expressing MyD88-mNeonGreen (MyD88-mNG), enabling real-time visualization and quantification of individual Myddosome assemblies at high resolution (Fig. 4A).

**Figure 4.**
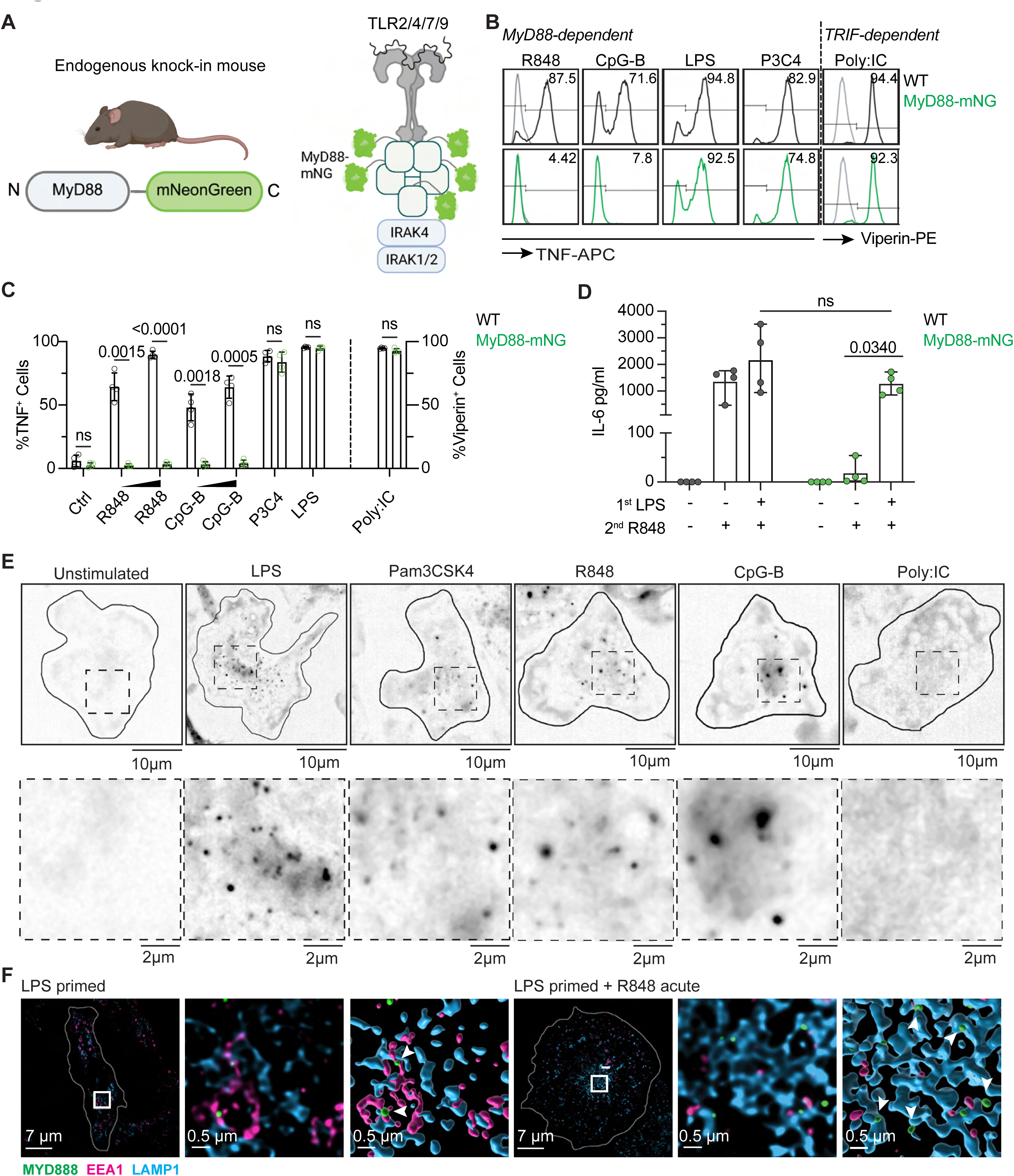
Real-time Myddosome visualization in stimulated macrophages. (**A)** Schematic of MyD88-mNG knock-in mouse and Myddosome assembly downstream of activated TLR2/4/7/9. **(B)** Intracellular TNF staining of WT and MyD88-mNG Hoxb8 macrophages in response to TLR stimulation for 6h: MyD88-dependent TLR4 (LPS 100ng/ml), TLR2 (Pam3CSK4 200ng/ml), TLR7 (R848 500ng/ml), TLR9 (CpG-B 1µM), and MyD88-independent TLR3 (Poly:IC 10µg/ml). Representative experiment out of N=4. **(C)** Pooled repeats of (B). P-values were calculated with a paired t-test. **(D)** IL-6 ELISA of WT and MyD88-mNG macrophages primed for 24h with 100ng/ml LPS and re-stimulated with R848 500ng/ml (N=4). P-values were calculated with two-way ANOVA with Tukey’s multiple comparisons test. **(E)** Widefield imaging of Myddosome formation in MyD88-mNG macrophages stimulated for 1 hour with the TLR ligands and concentrations as used in (B). **(F)** Fixed SIM microscopy of MyD88-mNG macrophages stimulated with LPS 100ng/ml for 1 hour or primed with LPS for 24h and then stimulated with R848 500ng/ml for 1 hour, co-stained with Lamp1 and EEA1. White arrows point to Myddosomes. ns: not significant.

First, we generated Hoxb8-immortalized progenitors from the bone marrow of WT and MyD88-mNG KI mice and differentiated them into macrophages. To assess MyD88-dependent signaling, these cells were stimulated for 6h with LPS (TLR4), Pam3CSK4 (TLR2), R848 (TLR7), CpG-B (TLR9), and Poly:IC (TLR3), and the cytokine output was compared to that of conventional bone marrow-derived macrophages (BMDMs) (Fig. 4B, C; Supp. Fig. 3A, B). Unexpectedly, whereas tagged MyD88 supported intact TLR4 signaling from the plasma membrane, TLR7 and 9 signaling from endosomal membranes was highly attenuated in both Hoxb8-derived macrophages and BMDMs derived from the reporter mouse. This observation suggests a fundamental difference in Myddosome formation at distinct subcellular locations, with endosomal membranes experiencing a limiting constraint in oligomerization of tagged MyD88. As expected, TRIF-dependent IFN-I production upon Poly:IC stimulation was unaffected, as evidenced by Viperin ICS staining (Fig. 4B). We next asked whether LPS priming could restore TLR7/9 activation in the attenuated MyD88-mNG cells. Strikingly, LPS priming fully rescued TLR7/9 responses to levels comparable to WT cells (Fig. 4D). These results demonstrated that 1) the LPS priming effect on TLR7 was preserved in MyD88-mNG KI macrophages, and, more importantly, that 2) LPS priming relieves a constraint on MyD88 signaling, enabling more efficient signal transduction from endosomal membranes even under suboptimal conditions. We therefore concluded that this reporter system provides a powerful tool for probing MyD88 signaling in both basal and primed states and can be used to analyze how LPS exposure affects Myddosome formation and contributes to TLR7 training.

We then characterized Myddosome puncta formation in MyD88-mNG macrophages in response to various TLR stimuli. Macrophages were stimulated for 90 minutes, and puncta formation was assessed in real-time using widefield microscopy. As expected, MyD88 puncta were observed in all MyD88-dependent conditions, but not upon stimulation with the MyD88-independent TLR3 agonist Poly:IC (Fig. 4E). To verify the identity of these puncta, we stably expressed IRAK2-mScarlet in Hoxb8-immortalized MyD88-mNG KI progenitors. Upon TLR7 stimulation, macrophages displayed co-localization of MyD88-mNG with IRAK2-mScarlet, confirming that the observed puncta represent active Myddosome complexes (Supp. Fig. 3C). To determine the subcellular localization of Myddosome formation, we stimulated TLR4 and TLR7 in MyD88-mNG Hoxb8 macrophages for 1.5 hours and imaged cells using structured illumination microscopy. At this time point, TLR4 had largely internalized from the plasma membrane and continued to signal from EEA1^+^ early endosomal compartments (Fig. 4F), as evidenced by colocalization with Myddosome puncta and consistent with previous reports (Ciesielska et al., 2021; Husebye et al., 2006). To overcome the signaling attenuation of TLR7 and being able to image TLR7 Myddosomes, we primed cells with LPS first. Upon subsequent TLR7 stimulation, MyD88 puncta colocalized almost exclusively with the late endosomal/lysosomal marker Lamp1 (Fig. 4F), the primary compartment from which TLR7 signals. For both TLRs, we also observed a substantial fraction of free-floating Myddosomes in the cytosol, not associated with endosomal vesicles. These were greater in numbers after LPS stimulation, but also visible after TLR7 stimulation following LPS priming (Supp. Figure 3F). These cytosolic structures may represent Myddosomes that have released from their receptor after initial seeding at the respective membrane and which have been proposed to contribute to the overall signaling activity of a given cell (Fisch et al., 2024). In summary, our findings demonstrate that the MyD88-mNG KI system allows spatially resolved analysis of signalosome formation at distinct subcellular compartments, providing a platform to dissect how LPS priming modulates compartment-specific TLR signaling.

### 5. LPS priming enhances Myddosome nucleation downstream of TLR7

If training is encoded at the level of signalosome nucleation, LPS-primed macrophages should exhibit enhanced Myddosome formation upon TLR7 restimulation. To quantify Myddosome assemblies after LPS priming, MyD88-mNG macrophages were left unstimulated or primed with LPS for 24h, re-stimulated with R848 and imaged in real-time. Consistent with attenuated TLR7 signaling in this system, puncta formation in R848-stimulated cells was low. Importantly, although initial LPS stimulation induced a substantial number of Myddosomes, most of these structures disappeared after 24 hours of priming, consistent with a return to baseline. In accordance with the observed TLR7 training effect, puncta formation was greatly enhanced in primed cells upon R848 restimulation (Figure 5A). LPS priming increased both the total number of Myddosomes per cell, with some cells containing up to 150 assemblies per cell (Fig. 5B), and the fraction of macrophages containing Myddosomes (Fig. 5C). In unprimed cells, approximately 40% of macrophages formed Myddosome puncta after R848 treatment, and at low numbers that did not translate into robust signaling outputs. This proportion increased up to 70% after LPS priming. (Fig. 5B, C). These findings demonstrate that LPS priming strongly amplifies Myddosome nucleation at endosomes. Conversely, during endotoxin tolerance, Myddosome formation is markedly attenuated during a second challenge with LPS (Fig. 5B, C). We confirmed that TLR7 signaling amplification was not due to upregulated MyD88 expression after LPS priming. (Figure 5D, E; Supp. Figure 3D, E). Proteome profiling further confirmed that expression of other core signaling molecules in the TLR pathway, including IRAK1, 2, 4, and RelA, remained unchanged. Therefore, we conclude enhanced TLR7 signaling as a result of LPS priming is driven by enhanced assembly of Myddosome complexes at endosomal membranes.

**Figure 5.**
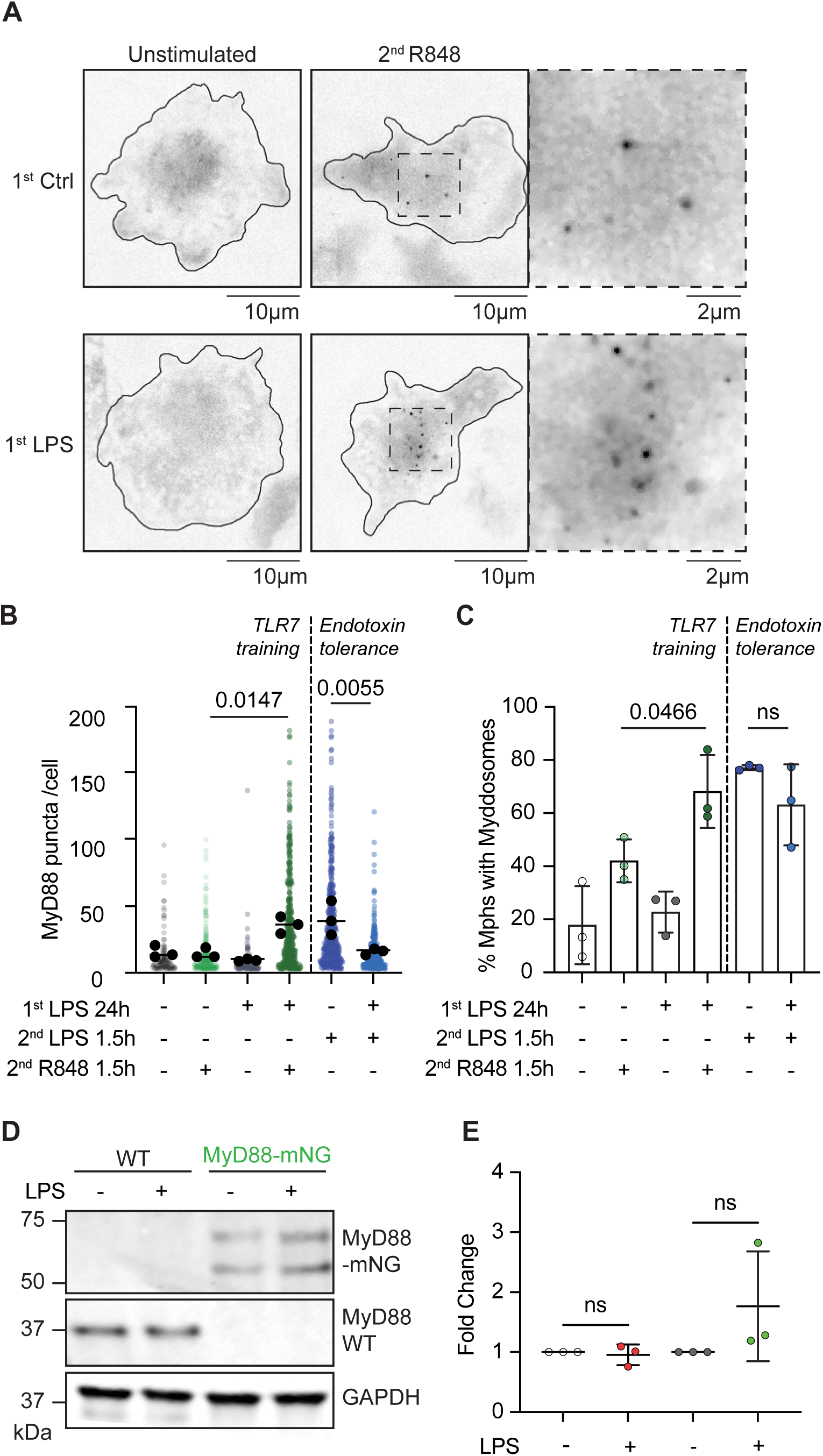
TLR7-dependent Myddosome assembly is enhanced in LPS-primed macrophages. **(A)** Representative images of widefield microscopy of TLR7-dependent Myddosome formations in MyD88-mNG macrophages with or without prior LPS priming. **(B)** Quantification of MyD88 puncta per cell (only cells that contained at least 3 Myddosomes were included). Graph shows all individual cells from 3 independent repeats and corresponding means (N=3). Total number of analyzed cells: unstimulated=652, R848 only=620, LPS+R848=550, LPS acute=565, LPS no secondary stim.=504, LPS+LPS restim.=523. **(C)** Quantification of the cell fraction containing at least 3 Myddosomes: data is pooled from three independent experiments (N=3). Total number of analyzed cells: unstimulated=99, R848 only=262, LPS+R848=369, LPS acute=437, LPS no secondary stim.=121, LPS+LPS restim.=303. P-values for (B) and (C) were calculated by unpaired t-test. **(D)** Western blot analysis of WT MyD88 and MyD88-mNeonGreen in naïve and LPS-primed macrophages. Representative blot out of N=3. **E:** Quantification of (D). P-values were calculated by unpaired t-test, ns: not significant.

In conclusion, we identify a novel mechanism of trained immunity in which LPS exposure alters the threshold of TLR7 signal transduction by enhancing Myddosome nucleation, thereby strongly increasing inflammatory cytokine outputs. The precise molecular mechanism of this amplification will require future investigation, but we speculate that priming-mediated changes in the protein or lipid composition of endosomes could account for the enhanced signaling capacity. Alternatively, residual Myddosomes that assembled in response to the initial LPS priming response might provide ready-to-use pre-assembled complexes that enhance faster and more efficient signalosome assembly upon TLR7 stimulation. These residual and preformed Myddosomes are possibly smaller in size to those we visualized after stimulation. They are most likely closer in size to the solved Myddosome structure and those observed after TLR4 or IL-1 stimulation (Deliz-Aguirre et al., 2021; Fisch et al., 2024) in some unstimulated cells (Moncrieffe et al., 2020) and therefore, may have escaped visualization using our Myd88-mNG reporter. Regardless of these uncertainties, a central conceptual advance of this work is the demonstration that cellular adaptation can be encoded not only through long-lasting epigenetic and metabolic changes, but also through rapid, receptor-proximal reprogramming of signal transduction.

Here, we demonstrate that cellular adaptation and the reprogramming of signaling pathways can occur in a compartment- and receptor-specific manner. Evolutionarily, such rapid adaptations may represent a conserved strategy for fine-tuning immune responses based on prior pathogen exposure during overlapping or sequential infections. However, this mechanism may carry immunopathological risks. Enhanced endosomal TLR sensitivity could lower the activation threshold for self-derived nucleic-acids and promote autoimmunity in genetically predisposed individuals. Epidemiological studies point to a clear association between infections and autoimmune disease, irrespective of pathogen type but with a strong temporal component (Nielsen et al., 2016). Our data provide a mechanistic framework, by which bacterial infections could precede or exacerbate autoimmune flares via cell-intrinsic signal adaptations. Understanding how immune cells reconfigure their intracellular signaling landscapes in response to environmental triggers may reveal opportunities for harnessing trained immunity in immunomodulatory therapies, such as vaccines or cancer treatments, and preventing its maladaptive consequences in chronic inflammation and autoimmunity.

---

**Figure S1.**
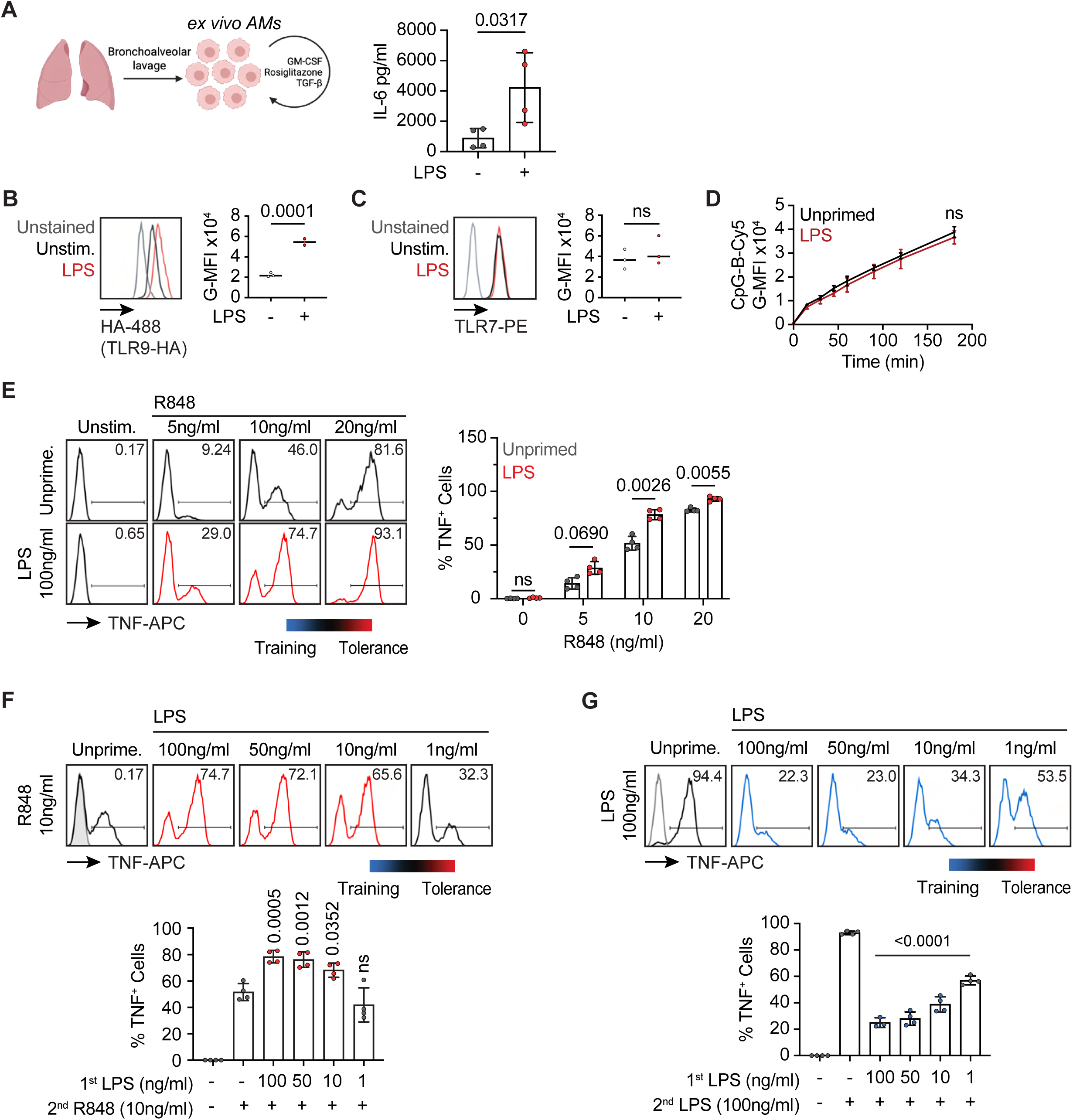
Characterization of trained TLR7 responses. **(A)** LPS primes TLR7-dependent IL-6 production in murine *ex vivo* alveolar macrophages (N=4). (**B-C)** TLR7 or TLR9-HA staining in Hoxb8 macrophages upon priming with LPS 100ng/ml (N=3). **(D)** Hoxb8 macrophages were primed with LPS 100ng/ml or left unprimed for 24h; cells were then fed with CpG-B-Cy5 for 0-180 minutes and uptake was quantified with flow cytometry (N=3). P values were calculated with ratio paired t-test at t=180min. **(E)** Intracellular TNF staining of Hoxb8 macrophages primed with LPS 100ng/ml and restimulated with increasing concentrations of R848: 5, 10, or 20ng/ml (N=4). P-values were calculated with two-way ANOVA and Tukey’s multiple comparison test **(F)** Intracellular TNF staining of Hoxb8 macrophages primed with decreasing concentrations of LPS: 100, 50, 10 or 1ng/ml. After 24h, cells were restimulated with 10ng/ml R848 (N=4). P-values were calculated with one-way ANOVA with Tukey’s post-test and each compared to the unprimed+R848 group. ns: not significant**G)** Hoxb8 macrophages were primed as in E) and then restimulated with 100ng/ml LPS. Flow histograms show a representative experiment out of N=4. Graphs show pooled repeats (N=4). P-values were calculated with one-way ANOVA with Tukey’s post-test. ns: not significant.

**Figure S2.**
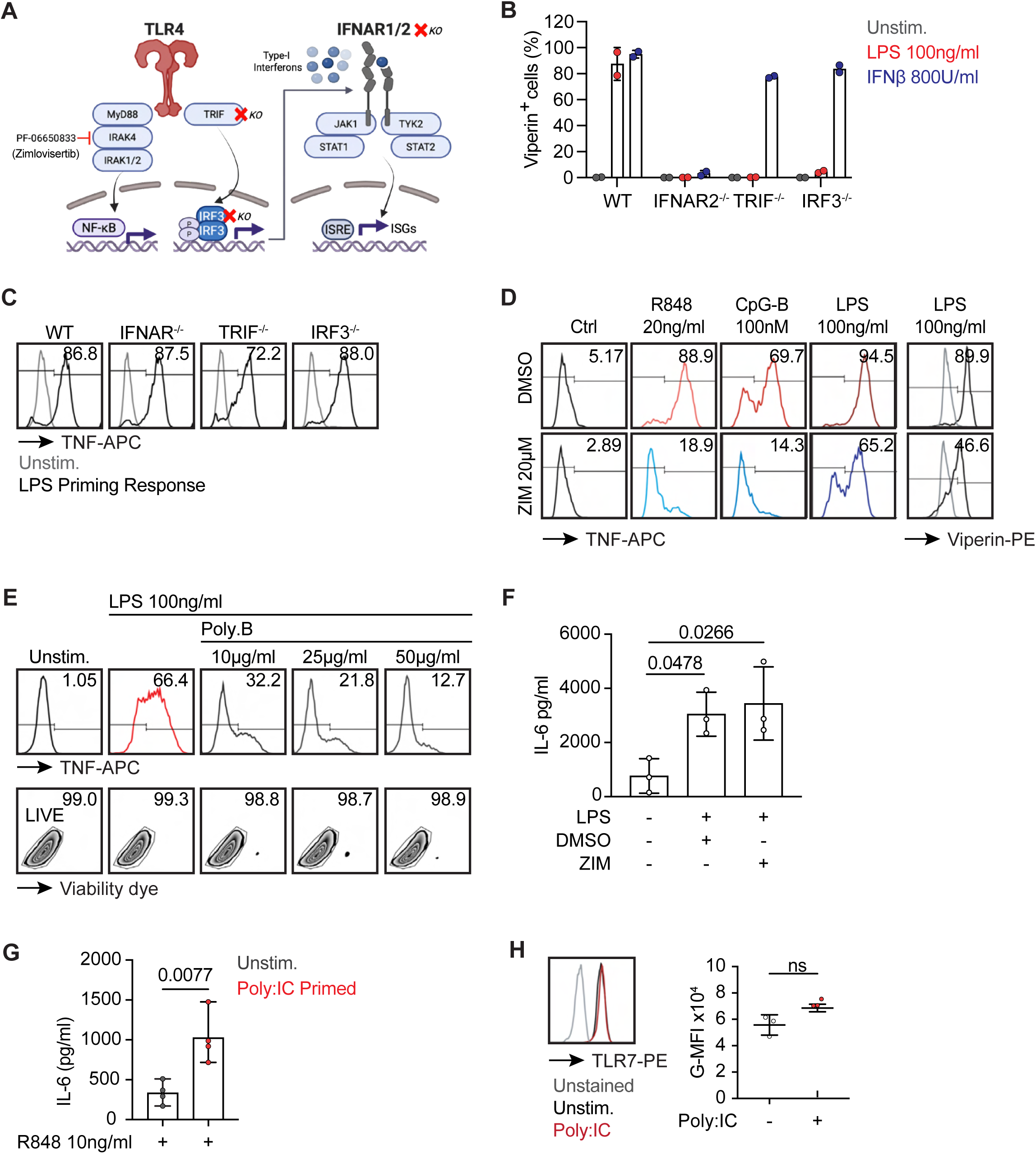
Knockout cell line validation and IRAK4 inhibition. **(A)** Schematic of TLR4-dependent signaling pathways and respective KOs and inhibitors used. **(B)** Functional validation of CRISPR-Cas9 knockouts of IFNAR, TRIF or IRF3 in Hoxb8 macrophages. Cells were stimulated with LPS 100ng/ml or IFN-β 800U/ml, and expression of the interferon stimulated gene Viperin was measured by intracellular staining and flow cytometry (each dot represents an independent KO line) **(C)** Intracellular TNF staining of knockout lines after 100ng/ml LPS stimulation for 6 hours (N=2). **(D)** Characterization of IRAK4 inhibitor Zimlovisertib: WT Hoxb8 macrophages were treated with Zimlovisertib (20µM) or vehicle and stimulated with TLR ligands: R848 20ng/ml, CpG-B 100nM, LPS 100ng/ml. Viperin and TNF production was measured by intracellular staining (N=1). **(E)** Intracellular TNF staining and viability control for Polymyxin-B: Hoxb8 WT macrophages were stimulated with LPS 100ng/ml for 6 hours after preincubation with increasing doses of Polymyxin-B (0, 10, 25, 50µg/ml) (N=1). **(F)** IL6-ELISA of WT Hoxb8 macrophages primed for 24 hours with LPS 100ng/ml in presence or absence of IRAK4 inhibitor Zimlovisertib (20µM) and re-stimulated with R848 10ng/ml or CpG-B 50nM (N=3) after washout of the inhibitor. **(G)** IL-6 ELISA of Hoxb8 macrophages primed with Poly:IC 10µg/ml for 24 hours and re-stimulated with R848 10ngml for 6 hours (N=4). **(H)** TLR7 staining of Hoxb8 macrophages left unprimed or primed with Poly:IC 10µg/ml for 24 hours. Representative histogram out of N=3. Graph shows pooled repeats. P-values were calculated with ratio paired t-test, ns: not significant.

**Figure S3:**
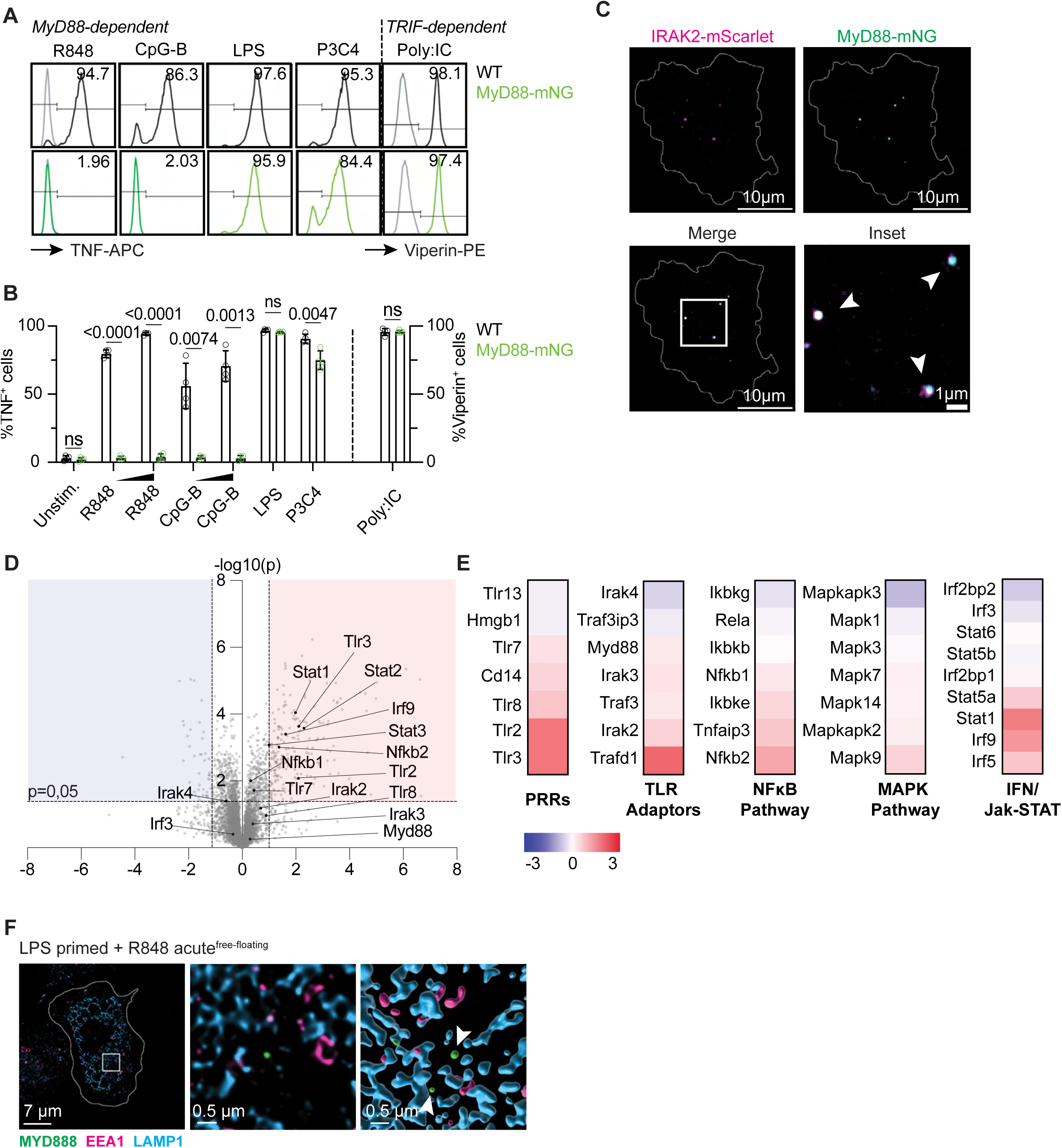
Characterization of TLR signaling in MyD88-mNG macrophages. **(A)** Intracellular TNF staining of WT and MyD88-mNG bone marrow-derived macrophages in response to TLR stimulation for 6h: MyD88-dependent TLR4 (LPS 100ng/ml), TLR2 (Pam3CSK4 200ng/ml), TLR7 (R848 500ng/ml), TLR9 (CpG-B 1µM), and MyD88-independent TLR3 (Poly:IC 10µg/ml). Representative histograms out of N=4. **(B)** Pooled repeats of (A). P-values were calculated with paired t-test. **(C)** Representative fixed SIM image of MyD88-mNG Hoxb8 macrophages stably expressing IRAK2-mScarlet and stimulated with R848 (500ng/ml) for 1 hour. **(D)** Mass spectrometry analysis of naïve and LPS primed whole cell lysates. Volcano blot shows log2 fold change over p-value of differentially expressed proteins (N=4). **(E)** Heat map shows log2fold change of mass spec. hits of relevant pathway proteins. Color code: Red = upregulated, blue = downregulated. (F) Free-floating Myddosomes: Fixed SIM microscopy of MyD88-mNG macrophages primed with LPS for 24h and then stimulated with R848 500ng/ml for 1 hour, co-stained with Lamp1 and EEA1. White arrows point to Myddosomes.

## Material and Methods

### Cell and tissue culture conditions

Myeloid Hoxb8 progenitors were cultured in RPMI 1640 medium conditioned with 2% granulocyte-monocyte colony-stimulating factor (GM-CSF), and supplemented with 10% FCS (Sigma-Aldrich #F0804), 1% L-Glutamine (Gibco #15030081), 1% sodium pyruvate (Gibco #11360070), 1% Hepes (Gibco #15630056), 1% penicillin-streptomycin (Gibco #15140122), 0.02% β-estradiol (Sigma-Aldrich #E8875) and 0,0003% β-mercaptoethanol (Gibco #21985-023). GM-CSF was produced by a B16 murine melanoma cell line.

Cas9 expressing murine Hoxb8 progenitors were a gift from the Barton Lab, Berkeley, California (USA). Hoxb8 progenitors expressing TLR9-HA were derived from the TLR9-HA-IRES-GFP KI mouse (Price et al., 2018).

Human embryonic kidney-293T (Hek293T) cells (American Type Culture Collection), and GP2-293 packaging cell line (Clontech) were cultured in Dulbecco’s modified eagle medium (DMEM) (Invitrogen #10938-025) supplemented with 10% FCS, 1% penicillin-streptomycin, 1% sodium pyruvate, 1% L-Glutamine, and 1% Hepes. All cell types were cultured at 37°C and 5% CO_2_.

### Generation of GM-CSF-derived Hoxb8 progenitors

For detailed protocol, refer to (Wang et al., 2006). In brief, Hoxb8 progenitors were derived from bone marrow of C57BL/6 mice. The bone marrow was flushed and cultured for 2 days in stem cell medium containing DMEM, 15% FCS, stem cell factor 25ng/ml (Peprotech) + IL-3 10ng/ml (Peprotech) + IL-6 20ng/ml (Peprotech). For immortalization, cells were spinfected with a retrovirus containing a FLAG-ER-Hoxb8-MSCV-Neo construct. Immortalized progenitors were selected by survival in progenitor medium containing β-estradiol and passaged for several days. Stocks were frozen and stored in liquid nitrogen.

### Differentiation of Hoxb8 macrophages

For macrophage differentiation, immortalized GM-CSF Hoxb8 progenitors were washed 2x in PBS and differentiated in macrophage medium (RPMI1640, 10% FCS, 1% L-glutamine, 1% sodium pyruvate, 1% Hepes, 1% Penicillin-Streptomycin, 10% conditioned M-CSF medium produced by a 3T3 mouse embryonic fibroblast cell line, and 0.0003% β-mercaptoethanol) in non-TC treated petri dishes for 7 days. The medium was refreshed on day 3. For experiments, macrophages were detached with cold PBS and scraped.

### Macrophage priming and stimulation procedure

Cells were replated in 96-well plates at 7x10^5^ cells per well and primed with LPS-B5 100ng/ml for 24h. The priming stimulation was then washed out two times with PBS. For cytokine readouts, cells were then incubated with the secondary ligand: R848 10ng/ml (TLR7), CpG-B 50nM (TLR9), Poly:IC 10µg/ml (TLR3), Pam3CSK4 200ng/ml (TLR2), or LPS-B5 100ng/ml (TLR4). For SIM microscopy of MyD88-mNG cells, higher ligand concentrations were used for TLR7 and TLR9: R848 500ng/ml and CpG-B 1µM.

### Culture and stimulation of ex vivo alveolar macrophages

*Ex vivo* alveolar macrophages (AMs) were isolated by bronchoalveolar lavage and were a kind gift by Bastian Opitz, Charité, Berlin. AM isolation and culture was performed according to (Busch et al., 2019). In brief, cells were maintained at a density of 3-4x10^5^ cells per well in TC-treated 6-well plates in complete medium consisting of RPMI1640, 10% FCS, 1% L-glutamine, 1% sodium pyruvate, 1% Hepes, 1% Penicillin-Streptomycin supplemented with 30ng/ml recombinant murine GM-CSF (Peprotech #315-03), 10ng/ml human TGFβ (Peprotech #100-21), and 1µM Rosiglitazone (Sigma Aldrich #R2408). Cells were incubated at 37°C and 5% CO2. Cells were fed every 2nd day and split upon confluency. For stimulation, AMs were detached with EDTA-Trypsin and replated at 7x10^5^ cells per well in 96-well plates and left to settle for 24 hours. The next day, cells were primed and restimulated as indicated.

### CRISPR-Cas9 knockout in Hoxb8 macrophages

For a detailed protocol, refer to (Roberts et al., 2019). In brief, Cas9-Hoxb8 progenitors were transduced via spinfection with lentivirus containing the pLentiGuidePuro (Addgene #52963) expressing the respective small guide RNA (sgRNA) sequence, which had been cloned using the BsmBI restriction enzyme site. sgRNA sequences used in this study are listed in key resources table. Cells were put on Puromycin selection 48h after spinfection.

### Stable expression of IRAK-4-mScarlet in Hoxb8 macrophages

For the production of lentiviral particles, 5x10^5^ human embryonic kidney (HEK) 293T cells/well were plated in a 6-well plate in complete DMEM at 37°C and 5% CO_2_ one day before transfection. The following day, pHR-dSV-IRAK2-mScarlet, envelope plasmid pVSV-G, and packaging plasmid pPAX2 were transfected using Lipofectamine 3000 reagent according to the manufacturer’s instructions. 10 hours after transfection, the media was exchanged with progenitor medium without β-estradiol. One day post transfection, the packaging cells were moved to 32°C and 5% CO_2_. Two days post transfection, viral supernatant was collected, filtered through a 0.45μm syringe filter, and supplemented with 0,02% β-estradiol and 5μg/ml polybrene. In a 6-well plate, 5x10^5^ Hoxb8-derived MyD88-mNeonGreen progenitors per well (in 0.5ml) were plated and the viral supernatant (2ml) added. For transduction, cells were spinfected for 30 min at 1000g and 32°C and incubated overnight at 32°C and 5% CO_2_. Successfully transduced cells were selected with 5μg/ml puromycin (A11138-03, Gibco) starting 48h post transduction. About one week later, cells were FACS sorted to select mNeonGreen/mScarlet double-positive cells.

### Western Blot analysis

For immunoblots, 1.2x10^6^ cells were lysed in a lysis buffer containing 50mM TRIS-HCl, 150mM NaCl, 5mM EDTA, 1% NP-40, 1x EDTA-free Protease Inhibitor tablet (Roche, #11873580001). For phospho-blots, lysis buffer also contained 1x PhosSTOP phosphatase inhibitor tablet (Roche, #04906 837001). Cells were lysed in 100µl lysis buffer on ice for 1h. Lysates were then centrifuged at maximum speed for 25 minutes at 4°C. Clear supernatant was transferred to a new tube, and 4X Laemmli Sample buffer containing DTT (Bio-Rad #1610747) was added for a final concentration of 1X. Samples were left to denature at room temperature for 1h and transferred to -20°C until further use. For SDS-PAGE analysis, samples were applied to 15-well, 15µl, 4-15% polyacrylamide gels (Bio-Rad Mini PROTEAN Precast Gels #4561086) and transferred to Immobilon polyvinylidene difluoride membranes (Millipore) in a Trans-Blot Turbo transfer system (Bio-Rad). Membranes were blocked with TBS+BSA 5% or protein-free Li-cor blocking buffer (Li-cor Biosciences #92780001). Primary antibodies were diluted in blocking buffer + Sodium-Azide, secondary antibodies in blocking buffer only. Blot development occurred as indicated with either ECL blotting substrate (Thermo Scientific #11527271) on the ChemiDoc imaging system (Bio-Rad) or secondary fluorescent antibodies (AlexaFluor680) for detection with the Li-cor scanning system for quantitative analysis. Band intensities were quantified using FIJI/Image software. Images were imported as TIFFs, and densitometry was performed using the Gel Analysis workflow. Briefly, rectangular regions of interest of identical size were drawn around each band, and the intensity profiles were generated. The wand tool was used to measure the area under each peak, corresponding to band intensity. Density values were normalized to the corresponding loading control.

### Flow Cytometry

Cells were seeded into non-TC treated U-bottom 96-well plates at 7x10^5^ cells per well and left to settle for 24 hours. Cells were then left unstimulated or stimulated with ligands as indicated. For intracellular TNF stainings, Brefeldin A (GolgiPlug, BD Biosciences #555029, 1:1000) was added to each well 20 minutes after stimulation. Cells were collected after different time points as indicated, and dead cells were stained with fixable live/dead dye eFluor506 in PBS. Cells were then fixed with the BD Cytofix/Cytoperm kit (#554714) according to the manufacturer’s instructions. To block Fc receptors, cells were treated with anti-mouse CD16/CD32 (1:100, Biolegend #156603) for 10 minutes and then stained for TLRs or cytokines for 1 hour (antibodies and concentrations indicated in extended materials). Data was acquired on the CytoFLEX machine (Beckman Coulter) and analysed with FlowJo software.

### IFNs-I bioassay

L292-ISRE reporter cells were cultured in complete RPMI-1640 medium and plated at 2x10^4^ cells per well in 100µl in 96-well TC-treated plates one day prior to the assay. For the IFNβ standard curve, recombinant mouse IFNβ (Biolegend, #581302) was diluted to a top standard concentration of 8ng/ml. A 2-fold serial dilution was performed 10 times to generate a standard curve range from 8ng/ml to 7.8pg/ml. Medium was removed from L292-ISRE cells and 100µl of supernatants from experimental stimulations or standard dilutions were added. Cells were incubated overnight at 37°C. After incubation, the medium was removed and 50µl of 1X passive lysis buffer (Promega) was added to each well. Plates were incubated shaking for 15 minutes at room temperature. 20µl of the lysate were added to a new white 96-well plate, and luciferase activity was measured using the Luciferase Assay System (Promega). Luminescence was recorded for 10 seconds per well following automatic injection of 50µl luciferase substrate. Luminescence values from the standard curve were used to interpolate experimental samples.

### CpG-B uptake assay

For ligand uptake quantification, Hoxb8 WT macrophages were plated at 7x10^5^ cells/well in non-TC treated 96-well plates. Cells were left unprimed or stimulated with LPS-B5 100ng/ml for 24 hours. LPS was washed off two times with PBS, and cells were incubated with CpG-B-Cy5 (1:2000) for durations indicated. Cy5 intensity was measured by flow cytometry as described above and analyzed using FlowJo.

### Immunofluorescence

Macrophages were plated onto (no. 1.5H) glass coverslips at a density of 2.5 × 10⁵ cells per coverslip and allowed to settle for 24 hours. Where indicated, cells were primed with 100 ng/ml LPS-B5. The following day, cells were rinsed with PBS and stimulated as described. After 1.5 hours, samples were washed once with PBS and fixed in glyoxal solution (Richter et al., 2018) for 15 min at room temperature. Macrophages were then permeabilized with 0.5% saponin for 5 min, followed by a 10-min incubation in 50 mM ammonium chloride/0.1% saponin in PBS to reduce PFA autofluorescence. Between each indicated step, cells were rinsed with PBS. Afterward, samples were blocked for 1 h in 3% BSA/0.1% saponin/4% horse serum (Gibco #26050070) in PBS and incubated overnight at 4°C with primary antibodies diluted in 1% BSA/0.1% saponin in PBS. The next day, samples were washed three times with PBS and incubated for 1 hour at room temperature with secondary antibodies in blocking solution. Coverslips were washed three times with PBS and mounted onto glass slides using ProLong Glass (Invitrogen #P36982). Slides were cured for 24 h at room temperature and imaged on a Zeiss Elyra 7 with lattice structured illumination microscopy (SIM).

### Super-resolution structured illumination microscopy (SIM)

Microscopy was performed using a Zeiss Elyra 7 Lattice SIM microscope equipped with 405-, 488-, 561-, and 642-nm lasers. Images were acquired with a Plan-Apochromat 63×/NA 1.6 oil immersion objective, and two pco.edge 4.2 sCMOS (scientific complementary metal-oxide semiconductor) cameras. Live imaging was conducted at 37°C and 5% CO_2_, fixed-sample imaging at 30°C. To minimize acquisition time, live-cell z-stacks were acquired in Leap mode. All raw images were SIM-processed using SIM² mode in Zeiss ZEN Black 3.0 software. Multi-color fixed-cell images were additionally channel-aligned as described previously (Mishra et al., 2024). Image visualization and postprocessing steps, including color and contrast adjustments, z-projection generation, and addition of cell outlines and scale bars, were carried out in ImageJ/Fiji. Three-dimensional renderings of z-stacks were generated in Imaris 9.6.0.

### Generation of MyD88-mNeonGreen knock-in mice

The MyD88-mNeongreen knock-in (KI) mice were generated by inserting the mNeongreen coding sequence before the stop codon of *MyD88* at the endogenous locus using CRISPR/Cas9 editing technology. Briefly, specific sgRNAs targeting the region near the *MyD88* stop codon were designed using the Geneious software and the CRISPOR design webtool (http://crispor.tefor.net/). Guide RNA (GGCAAGGCGGGTCCAGAACC) was ordered as crRNA from IDT (Integrated DNA Technologies). crRNA was duplexed with tracrRNA (IDT Cat.no. 1072534) by mixing equimolar amounts. A double-stranded (ds) DNA donor template (ca. 1.7 kb) containing the mNeongreen fusion sequence was separated from the C-terminus of MyD88 by a 9 amino acid Glycine Serine linker (3xGGS). Mouse embryonic stem cells were transfected with RNP (44 pmol cr:tracr duplex and 37 pmol Cas9, IDT Cat.no. 1081061) and dsDNA donor template (500 ng/µl) using the NEON transfection system (settings: 1200 V, 20 ms, 2 pulses). Positive mES cell clones were identified and genotyped to confirm the in-frame fusion of mNeongreen with MyD88. Positive mESC clones were injected into 8-cell C57BL/6NCrl mouse embryos and transferred to the oviduct of pseudo-pregnant recipients. Male chimeras were confirmed by short tandem repeat analysis on sperm to ensure germline transmission. IVF was performed with positive sperm, and heterozygous animals were bred to homozygosity.

C57BL/NCrl mice were used to generate MyD88-mNeongreen KI animals. The generation of the MyD88-mNeongreen mouse was performed at the Genome Engineering and Transgenic Core Facility at the Max-Planck-Institute for Molecular Cell Biology and Genetics in Dresden and approved by the Saxony state authorities. Mouse breeding and the euthanasia of WT and KI mice lines for the harvesting of bone marrow progenitor cells were approved by the Berlin state authority Landesamt für Gesundheit und Soziales. Mice were bred at the Max-Planck-Institute for Infection Biology in Berlin, housed under specific pathogen-free conditions on a 12-hour light/dark cycle, and fed ad libitum.

### Live SIM microscopy and quantification of Myddosome assembly

For live SIM imaging of MyD88-mNeonGreen Hoxb8-derived macrophages, cells were seeded one day before imaging onto glass-bottom 8-well live-cell imaging chambers (Nunc Lab-Tek #155409) at 7 × 10⁴ cells per well in macrophage medium. Cells were either primed with 100 ng/ml LPS-B5 or left untreated. After 24 hours, the priming stimulus was removed, and cells were rinsed once with PBS. Macrophages were then re-stimulated with 500 ng/ml R848, 100 ng/ml LPS-B5, or left unstimulated in macrophage medium prepared with phenol red–free RPMI (Gibco #11835030). Images were acquired 1 hour after stimulation. Raw z-stacks were SIM-processed, and corresponding laser widefield (WF) images were reconstructed using Zeiss ZEN Black software.

To quantify Myddosome assemblies, SIM-processed and reconstructed laser WF z-stacks were merged into maximum-intensity projections and exported as TIFF files using ImageJ/Fiji. Using CellProfiler 4.2.7 (Carpenter et al., 2006), laser WF maximum-projection images were median-blurred, and an Otsu threshold was applied to segment individual cells. For improved puncta identification, cells were classified into five groups based on their mean MyD88-mNeonGreen fluorescence intensity. A noise-subtraction step was applied to SIM maximum-projection images to enhance puncta structures. MyD88 puncta were then segmented using an adaptive Otsu thresholding approach, with thresholds settings individually optimized for each intensity group. Following segmentation, Myddosomes were associated with their respective cell objects, and the number of Myddosomes per cell was quantified. Values were exported, and the percentage of cells containing MyD88 puncta was calculated in R (4.3.2). Macrophages were classified as Myddosome-positive if they contained ≥3 puncta. Myddosome counts >200 were excluded because they reflected segmentation errors.

Images acquired in widefield mode were subjected to a rolling-ball background subtraction (radius = 100 pixels), followed by a median filter (radius = 2 pixels) using ImageJ/Fiji to enhance Myddosome puncta.

### Proteome profiling

PBS-washed macrophage cell pellets derived from Hoxb8-immortalized bone marrow progenitors of 12 mice were lysed under denaturing conditions in 300 µL of a buffer containing 3 M guanidinium chloride (GdmCl), 10 mM tris(2-carboxyethyl)phosphine (TCEP), 40 mM chloroacetamide, and 100 mM Tris-HCl, pH 8.5. Lysates were denatured at 95°C for 10 min, shaking at 1000 rpm in a thermal shaker, and sonicated in a water bath for 10 min. The protein concentration of each sample was measured with a BCA protein assay kit (23252, Thermo Scientific, USA). 500 ng protein was used per sample and diluted with a dilution buffer containing 10% acetonitrile and 25 mM Tris-HCl, pH 8.0, to reach a 1 M GdmCl concentration. Then, proteins were digested with LysC (Roche, Basel, Switzerland; enzyme-to-protein ratio 1:50, MS-grade) by shaking at 800 rpm at 37°C for 3 hours. The digestion mixture was diluted again with the same dilution buffer to reach 0.5 M GdmCl, followed by tryptic digestion (Roche, enzyme to protein ratio 1:50, MS-grade) and incubation at 37°C overnight in a thermal shaker at 800 rpm. Peptides were acidified with formic acid to a final concentration of 2%, and the digests were loaded onto Evotip Pure (Evosep, Odense, Denmark) tips according to the manufacturer’s protocol. Peptide separation was carried out using nanoflow reverse-phase liquid chromatography (Evosep One, Evosep) with the Aurora Elite column (15 cm x 75 µm ID, C18 1.7 µm beads, IonOpticks, Victoria, Australia) and the 40-sample-per-day method (Whisper 40-SPD). LC-MS/MS was performed by using the data-independent acquisition (DIA) method with parallel accumulation serial fragmentation (PASEF). MS data were processed with Dia-NN (v1.8.2 beta 22) and searched against an in silico predicted mouse spectral library. The "match between run" feature was used, and the mass search range was set to m/z 400 to 1000.

MS data were processed with Perseus software. Differentially abundant proteins in LPS-primed cells were identified using a significance cutoff of p<0.05. For visualization, log2 fold changes were plotted against -log10(p) values to generate a volcano plot. Proteins associated with TLR signalling pathway were selected for focused analysis and visualized as a heatmap. All plots were generated using GraphPad Prism. The mass spectrometry proteomics data have been deposited to the ProteomeXchange Consortium via the PRIDE (Perez-Riverol et al., 2025) partner repository with the dataset identifier PXD070363.

### Statistics

Statistical parameters are reported in the figures and figure legends, wherein “n” refers to the number of replicates within the same experiment, and “N” refers to the number of independent experimental repeats. Representative data has been at least repeated three times and variances are shown as SD, unless otherwise stated in the figure legend. For comparison of two groups, paired or unpaired two-tailed t-test was used. Paired t-test was used to compare groups from pooled, independent experiments, with paired values coming from the same experiment.

Ratio paired t-test was used to compare two normalized groups. To compare means of more than two groups, one-way analysis of variance (ANOVA) was performed followed by Tukey’s multiple comparisons test. To compare two groups over a dose response, two-way ANOVA was performed followed by a Tukey’s multiple comparisons test. Data was considered statistically significant if p<0.05. Statistical analysis was performed using GraphPad Prism 9 Software (GaphPad Software Inc.).

## Data Availability

The mass spectrometry proteomics data have been deposited to the ProteomeXchange Consortium via the PRIDE^17^ partner repository with the dataset identifier PXD070363. Uncropped Western blots are provided as Source Data. All other data needed to support the conclusions of the paper are available in the main text or the supplementary materials and methods. Requests for key reagents, such as cell lines or plasmids, should be directed to Olivia Majer. These will be shared upon request, accompanied by a Material Transfer Agreement (MTA), for non-commercial research purposes. Request regarding the MyD88-mNG KI mouse should be directed to Marcus Taylor.

## Acknowledgements

We thank Bastian Opitz from the Charite, Berlin for donation of alveolar macrophages. We thank Louise Hentschel, Joy Lewis and Marilena Trigka for assistance with experiments. Schematics were created with BioRender.

## Author contributions

Conceptualization: AD, OM; Methodology: AD, FB, NQ, OM; Investigation: AD, FB, NQ, CH, JM, OT, DM, OM; Visualization: AD, FB, NQ, CH, OM; Funding acquisition: OM, Data curation (proteomics): DM; Project administration: OM, Supervision: OM, MT; Writing-original draft: AD, OM, MT; Writing – review & editing: AD, FB, NQ, CH, JM, OT, DM, OM.

## Supplementary Materials and Methods

### Antibodies and reagents

#### Anti-mouse antibodies and dyes for intracellular flow cytometry stainings

TruStain FcX^TM^ anti-CD16/32 Fc-Block (1:200, Biolegend #156603), anti-TNFɑ APC (1:200, Biolegend #506307), anti-HA-Tag AF488 (1:50, Cell Signaling Technology #2350S), anti-Viperin PE (1:100, BD Pharmingen^TM^ #565196), anti-TLR7 PE (1:200, Biolegend #160003), Live/Dead Dye eFluor 506 (1:500, Invitrogen #65-0866-14)

### Anti-mouse primary antibodies for western blotting

anti-GAPDH (1:1000, Invitrogen #MA5-15738), anti-TLR7 (1:500, R&D Systems #MAB7156), anti-HA (1:1000, Roche #3F10), anti-phospho-p38 (1:1000, Cell Signaling #4511T), anti-phospho p44/42 (ERK1/2; 1:1000, Cell Signaling #4377T), anti-p38 (1:1000, Cell Signaling #8690T), anti-p44/42 (ERK1/2; 1:1000, Cell Signaling #137F5), anti-MyD88 (1:1000, R&D Systems #AF3109).

### Secondary antibodies for western blotting

AlexaFluor 680 anti-mouse IgG (1:20.000, Invitrogen #A21057), AlexaFluor 680 anti-rabbit IgG (1:20.000, Invitrogen #), anti-mouse HRP-conjugated (1:20.0000, Jackson ImmunoResearch #111-035-166), anti-rabbit HRP-conjugated (1:20.000, Jackson ImmunoResearch #111-035-144)

### Anti-mouse primary antibodies for Immunofluorescence

anti-Lamp1 (1:1250 R&D Systems #AF4320), anti-EEA1 (1:800, Cell Signaling #3288S)

### Secondary antibodies for Immunofluorescence

AlexaFluor 647 anti-goat IgG (1:1250, Invitrogen #A32849), AlexaFluor 488 anti-mouse IgG (1:1000, Invitrogen #A32723), AlexaFluor 568 anti-rabbit IgG (1:1000, Invitrogen #A11011)

### TLR ligands

For priming experiments, TLR4 was stimulated with LPS-B5 from E.coli O55:B5 (Invivogen, #tlrl-pb5lps, ultrapure). TLR7 was stimulated with R848 (Resiquimod) (Invivogen, #tlrl-r848-1). TLR9 was stimulated with CpG-B DNA (ODN1668: TCCATGACGTTCCTGATGCT), synthesized by Integrated DNA Technologies. TLR3 was stimulated with Poly:IC (high molecular weight, Invivogen #tlrl-pic). Poly:IC was heated at 70°C for 10 minutes before stimulation. TLR2 was stimulated with Pam3CSK4 (Invivogen, #tlrl-pms).

### Chemicals and Inhibitors

For Polymxin-B neutralization of LPS, LPS ligand dilutions or cell culture supernatants were pre-incubated with Polymyxin-B (50µg/ml; Invivogen) for 15-30 minutes at room temperature before addition to naïve cells. Priming readouts were then performed by flow cytometry or ELISA as described above.

For IRAK4 inhibition with Zimlovisertib (PF-06650833), cells were pre-incubated with the inhibitor (20µM) for 1h before LPS priming. The inhibitor was then present throughout the entire 24h hours of priming. Cells were then washed 2x with PBS and rested for 2 hours to ensure no interference of the inhibitor with the secondary stimulation. Cells were then restimulated for flow cytometry or ELISA readouts as described above.

### Glyoxal solution preparation

Glyoxal solution (∼4 ml) was prepared by combining 2.835 ml ddH₂O, 0.789 ml absolute ethanol (for analysis), 0.313 ml glyoxal (∼40% stock solution; Merck #50649), and 0.03 ml acetic acid. The mixture was vortexed thoroughly, and the pH was adjusted to 4.5 using 1 M NaOH (Richter et al., 2018).

### Cytokine quantification

For cytokine analysis, cells were plated in non-TC treated flat bottom 96-well plates and left to settle for 24h. Cells were then stimulated with TLR ligands as indicated. Supernatants were harvested after 6h and stored at -20°C until further use. IL-6 was detected with Biolegend’s ELISA MAX Deluxe kit (#431304). Multiplex analysis of cytokines and chemokines was performed with Biolegend’s Mouse Anti-Virus Reponse Panel (#740622). Kits were used according to the manufacturer’s instructions.

### Key resources table

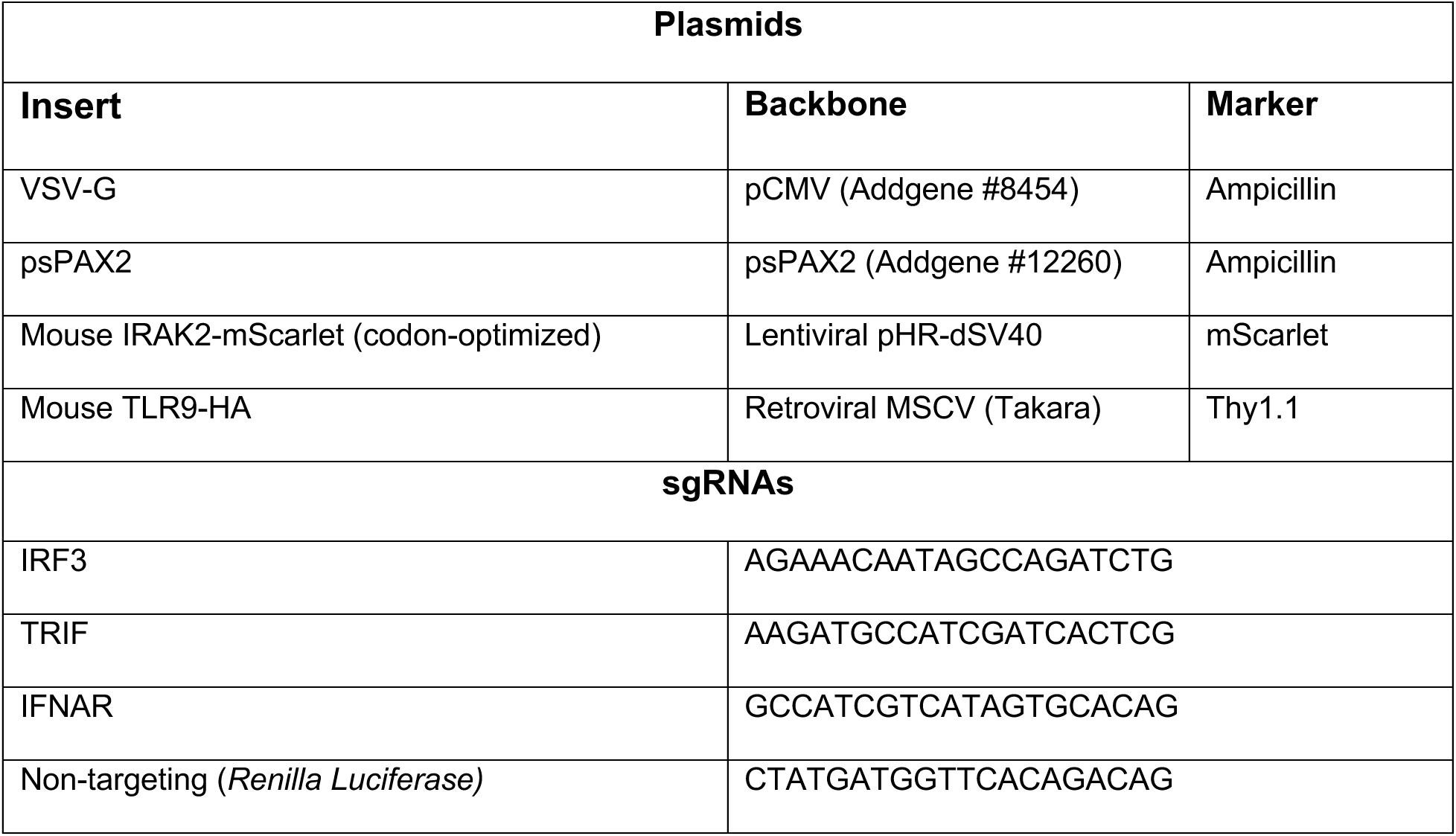

## References

Battaglia, M., and L.A. Garrett-Sinha. 2021. Bacterial infections in lupus: Roles in promoting immune activation and in pathogenesis of the disease. Journal of Translational Autoimmunity. 4:100078.

Brown, G.J., P.F. Cañete, H. Wang, A. Medhavy, J. Bones, J.A. Roco, Y. He, Y. Qin, J. Cappello, J.I. Ellyard, K. Bassett, Q. Shen, G. Burgio, Y. Zhang, C. Turnbull, X. Meng, P. Wu, E. Cho, L.A. Miosge, T.D. Andrews, M.A. Field, D. Tvorogov, A.F. Lopez, J.J. Babon, C.A. López, Á. Gónzalez-Murillo, D.C. Garulo, V. Pascual, T. Levy, E.J. Mallack, D.G. Calame, T. Lotze, J.R. Lupski, H. Ding, T.R. Ullah, G.D. Walters, M.E. Koina, M.C. Cook, N. Shen, C. De Lucas Collantes, B. Corry, M.P. Gantier, V. Athanasopoulos, and C.G. Vinuesa. 2022. TLR7 gain-of-function genetic variation causes human lupus. Nature. 605:349–356.

Busch, C.J., J. Favret, L. Geirsdottir, K. Molawi, and M.H. Sieweke. 2019. Isolation and Long-term Cultivation of Mouse Alveolar Macrophages. Bio Protoc. 9.

Carpenter, A.E., T.R. Jones, M.R. Lamprecht, C. Clarke, I.H. Kang, O. Friman, D.A. Guertin, J.H. Chang, R.A. Lindquist, J. Moffat, P. Golland, and D.M. Sabatini. 2006. CellProfiler: image analysis software for identifying and quantifying cell phenotypes. Genome Biol. 7:R100.

Ciesielska, A., M. Matyjek, and K. Kwiatkowska. 2021. TLR4 and CD14 trafficking and its influence on LPS-induced pro-inflammatory signaling. Cell Mol Life Sci. 78:1233–1261.

Dalpke, A.H., M.D. Lehner, T. Hartung, and K. Heeg. 2005. Differential effects of CpG-DNA in Toll-like receptor-2/-4/-9 tolerance and cross-tolerance. Immunology. 116:203–212.

Davies, K.J.A. 2016. Adaptive homeostasis. Molecular Aspects of Medicine. 49:1–7.

Davies, L.C., and P.R. Taylor. 2015. Tissue-resident macrophages: then and now. Immunology. 144:541–548.

De Nardo, D., K.R. Balka, Y. Cardona Gloria, V.R. Rao, E. Latz, and S.L. Masters. 2018. Interleukin-1 receptor–associated kinase 4 (IRAK4) plays a dual role in myddosome formation and Toll-like receptor signaling. Journal of Biological Chemistry. 293:15195–15207.

Deane, J.A., P. Pisitkun, R.S. Barrett, L. Feigenbaum, T. Town, J.M. Ward, R.A. Flavell, and S. Bolland. 2007. Control of toll-like receptor 7 expression is essential to restrict autoimmunity and dendritic cell proliferation. Immunity. 27:801–810.

Deliz-Aguirre, R., F. Cao, F.H.U. Gerpott, N. Auevechanichkul, M. Chupanova, Y. Mun, E. Ziska, and M.J. Taylor. 2021. MyD88 oligomer size functions as a physical threshold to trigger IL1R Myddosome signaling. J Cell Biol. 220.

Divangahi, M., P. Aaby, S.A. Khader, L.B. Barreiro, S. Bekkering, T. Chavakis, R. van Crevel, N. Curtis, A.R. DiNardo, J. Dominguez-Andres, R. Duivenvoorden, S. Fanucchi, Z. Fayad, E. Fuchs, M. Hamon, K.L. Jeffrey, N. Khan, L.A.B. Joosten, E. Kaufmann, E. Latz, G. Matarese, J.W.M. van der Meer, M. Mhlanga, S. Moorlag, W.J.M. Mulder, S. Naik, B. Novakovic, L. O’Neill, J. Ochando, K. Ozato, N.P. Riksen, R. Sauerwein, E.R. Sherwood, A. Schlitzer, J.L. Schultze, M.H. Sieweke, C.S. Benn, H. Stunnenberg, J. Sun, F.L. van de Veerdonk, S. Weis, D.L. Williams, R. Xavier, and M.G. Netea. 2021. Trained immunity, tolerance, priming and differentiation: distinct immunological processes. Nat Immunol. 22:2–6.

Doria, A., M. Canova, M. Tonon, M. Zen, E. Rampudda, N. Bassi, F. Atzeni, S. Zampieri, and A. Ghirardello. 2008. Infections as triggers and complications of systemic lupus erythematosus. Autoimmunity Reviews. 8:24–28.

Fisch, D., T. Zhang, H. Sun, W. Ma, Y. Tan, S.P. Gygi, D.E. Higgins, and J.C. Kagan. 2024. Molecular definition of the endogenous Toll-like receptor signalling pathways. Nature. 631:635–644.

Fukui, R., S. Saitoh, A. Kanno, M. Onji, T. Shibata, A. Ito, M. Onji, M. Matsumoto, S. Akira, N. Yoshida, and K. Miyake. 2011. Unc93B1 restricts systemic lethal inflammation by orchestrating Toll-like receptor 7 and 9 trafficking. Immunity. 35:69–81.

Husebye, H., O. Halaas, H. Stenmark, G. Tunheim, O. Sandanger, B. Bogen, A. Brech, E. Latz, and T. Espevik. 2006. Endocytic pathways regulate Toll-like receptor 4 signaling and link innate and adaptive immunity. EMBO J. 25:683–692.

Kobayashi, K., L.D. Hernandez, J.E. Galan, C.A. Janeway, Jr., R. Medzhitov, and R.A. Flavell. 2002. IRAK-M is a negative regulator of Toll-like receptor signaling. Cell. 110:191–202.

Lajqi, T., N. Kostlin-Gille, R. Bauer, S.G. Zarogiannis, E. Lajqi, V. Ajeti, S. Dietz, S.A. Kranig, J. Ruhle, A. Demaj, J. Hebel, M. Bartosova, D. Frommhold, H. Hudalla, and C. Gille. 2023. Training vs. Tolerance: The Yin/Yang of the Innate Immune System. Biomedicines. 11.

Latty, S.L., J. Sakai, L. Hopkins, B. Verstak, T. Paramo, N.A. Berglund, E. Cammarota, P. Cicuta, N.J. Gay, P.J. Bond, D. Klenerman, and C.E. Bryant. 2018. Activation of Toll-like receptors nucleates assembly of the MyDDosome signaling hub. Elife. 7.

Lin, S.C., Y.C. Lo, and H. Wu. 2010. Helical assembly in the MyD88-IRAK4-IRAK2 complex in TLR/IL-1R signalling. Nature. 465:885–890.

Majer, O., B. Liu, L.S.M. Kreuk, N. Krogan, and G.M. Barton. 2019. UNC93B1 recruits syntenin-1 to dampen TLR7 signalling and prevent autoimmunity. Nature. 575:366–370.

Medvedev, A.E., K.M. Kopydlowski, and S.N. Vogel. 2000. Inhibition of lipopolysaccharide-induced signal transduction in endotoxin-tolerized mouse macrophages: dysregulation of cytokine, chemokine, and toll-like receptor 2 and 4 gene expression. J Immunol. 164:5564–5574.

Mishra, H., C. Schlack-Leigers, E.L. Lim, O. Thieck, T. Magg, J. Raedler, C. Wolf, C. Klein, H. Ewers, M.A. Lee-Kirsch, D. Meierhofer, F. Hauck, and O. Majer. 2024. Disrupted degradative sorting of TLR7 is associated with human lupus. Science Immunology. 9:eadi9575.

Moncrieffe, M.C., D. Bollschweiler, B. Li, P.A. Penczek, L. Hopkins, C.E. Bryant, D. Klenerman, and N.J. Gay. 2020. MyD88 Death-Domain Oligomerization Determines Myddosome Architecture: Implications for Toll-like Receptor Signaling. Structure. 28:281–289 e283.

Napolitani, G., A. Rinaldi, F. Bertoni, F. Sallusto, and A. Lanzavecchia. 2005. Selected Toll-like receptor agonist combinations synergistically trigger a T helper type 1-polarizing program in dendritic cells. Nat Immunol. 6:769–776.

Netea, M.G., J. Domínguez-Andrés, L.B. Barreiro, T. Chavakis, M. Divangahi, E. Fuchs, L.A.B. Joosten, J.W.M. van der Meer, M.M. Mhlanga, W.J.M. Mulder, N.P. Riksen, A. Schlitzer, J.L. Schultze, C. Stabell Benn, J.C. Sun, R.J. Xavier, and E. Latz. 2020. Defining trained immunity and its role in health and disease. Nature Reviews Immunology. 20:375–388.

Nielsen, P.R., T.W. Kragstrup, B.W. Deleuran, and M.E. Benros. 2016. Infections as risk factor for autoimmune diseases - A nationwide study. J Autoimmun. 74:176–181.

Perez-Riverol, Y., C. Bandla, Deepti J. Kundu, S. Kamatchinathan, J. Bai, S. Hewapathirana, Nithu S. John, A. Prakash, M. Walzer, S. Wang, and Juan A. Vizcaíno. 2025. The PRIDE database at 20 years: 2025 update. Nucleic Acids Research. 53:D543–D553.

Price, A.E., K. Shamardani, K.A. Lugo, J. Deguine, A.W. Roberts, B.L. Lee, and G.M. Barton. 2018. A Map of Toll-like Receptor Expression in the Intestinal Epithelium Reveals Distinct Spatial, Cell Type-Specific, and Temporal Patterns. Immunity. 49:560–575 e566.

Rael, V.E., J.A. Yano, J.P. Huizar, L.C. Slayden, M.A. Weiss, E.A. Turcotte, J.M. Terry, W.Q. Zuo, I. Thiffault, T. Pastinen, E.G. Farrow, J.L. Jenkins, M.L. Becker, S.C. Wong, A.M. Stevens, C. Otten, E.J. Allenspach, D.E. Bonner, J.A. Bernstein, M.T. Wheeler, R.A. Saxton, B. Liu, O. Majer, G.M. Barton, and U.D. Network. 2024. Large-scale mutational analysis identifies UNC93B1 variants that drive TLR-mediated autoimmunity in mice and humans. J Exp Med. 221.

Richter, K.N., N.H. Revelo, K.J. Seitz, M.S. Helm, D. Sarkar, R.S. Saleeb, E. D’Este, J. Eberle, E. Wagner, C. Vogl, D.F. Lazaro, F. Richter, J. Coy-Vergara, G. Coceano, E.S. Boyden, R.R. Duncan, S.W. Hell, M.A. Lauterbach, S.E. Lehnart, T. Moser, T.F. Outeiro, P. Rehling, B. Schwappach, I. Testa, B. Zapiec, and S.O. Rizzoli. 2018. Glyoxal as an alternative fixative to formaldehyde in immunostaining and super-resolution microscopy. EMBO J. 37:139–159.

Roberts, A.W., L.M. Popov, G. Mitchell, K.L. Ching, D.J. Licht, G. Golovkine, G.M. Barton, and J.S. Cox. 2019. Cas9+ conditionally-immortalized macrophages as a tool for bacterial pathogenesis and beyond. eLife. 8:e45957.

Seeley, J.J., and S. Ghosh. 2017. Molecular mechanisms of innate memory and tolerance to LPS. J Leukoc Biol. 101:107–119.

Stout, R.D., and J. Suttles. 2004. Functional plasticity of macrophages: reversible adaptation to changing microenvironments. Journal of Leukocyte Biology. 76:509–513.

Wang, G.G., K.R. Calvo, M.P. Pasillas, D.B. Sykes, H. Häcker, and M.P. Kamps. 2006. Quantitative production of macrophages or neutrophils ex vivo using conditional Hoxb8. Nature Methods. 3:287–293.

West, M.A., and W. Heagy. 2002. Endotoxin tolerance: A review:. Critical Care Medicine. 30:S64–S73.

Wolf, C., E.L. Lim, M. Mokhtari, B. Kind, A. Odainic, E. Lara-Villacanas, S. Koss, S. Mages, K. Menzel, K. Engel, G. Dückers, B. Bernbeck, D.T. Schneider, K. Siepermann, T. Niehues, C.C. Goetzke, P. Durek, K. Minden, T. Dörner, A. Stittrich, F. Szelinski, G.M. Guerra, M. Massoud, M. Bieringer, C.C. De Oliveira Mann, E. Beltrán, T. Kallinich, M.-F. Mashreghi, S.V. Schmidt, E. Latz, J. Klughammer, O. Majer, and M.A. Lee-Kirsch. 2024. UNC93B1 variants underlie TLR7-dependent autoimmunity. Science Immunology. 9:eadi9769.

Wynn, T.A., A. Chawla, and J.W. Pollard. 2013. Macrophage biology in development, homeostasis and disease. Nature. 496:445–455.

Xiong, Y., F. Qiu, W. Piao, C. Song, L.M. Wahl, and A.E. Medvedev. 2011. Endotoxin tolerance impairs IL-1 receptor-associated kinase (IRAK) 4 and TGF-beta-activated kinase 1 activation, K63-linked polyubiquitination and assembly of IRAK1, TNF receptor-associated factor 6, and IkappaB kinase gamma and increases A20 expression. J Biol Chem. 286:7905–7916.

Zahalka, S., P. Starkl, M.L. Watzenboeck, A. Farhat, M. Radhouani, F. Deckert, A. Hladik, K. Lakovits, F. Oberndorfer, C. Lassnig, B. Strobl, K. Klavins, M. Matsushita, D.E. Sanin, K.M. Grzes, E.J. Pearce, A.-D. Gorki, and S. Knapp. 2022. Trained immunity of alveolar macrophages requires metabolic rewiring and type 1 interferon signaling. Mucosal Immunology. 15:896–907.

