## Supplementary figures and images for "Microbial Priming Enhances TLR7-Driven Myddosome Assembly Dynamics"

### Figures

#### Figure 1

##### Fig. 1D

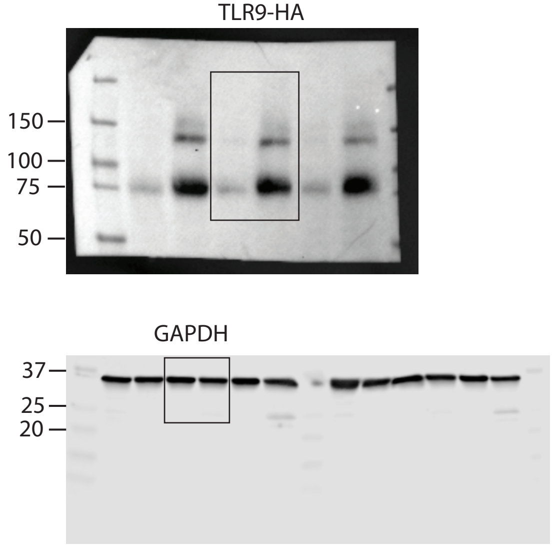


##### Fig. 1E

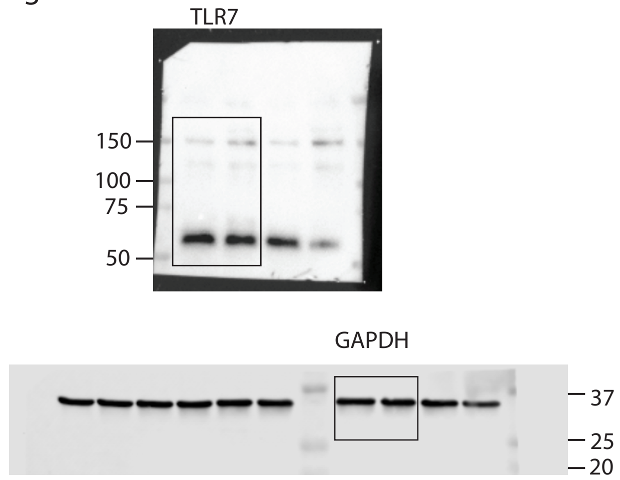


#### Figure 3

##### Fig. 3A

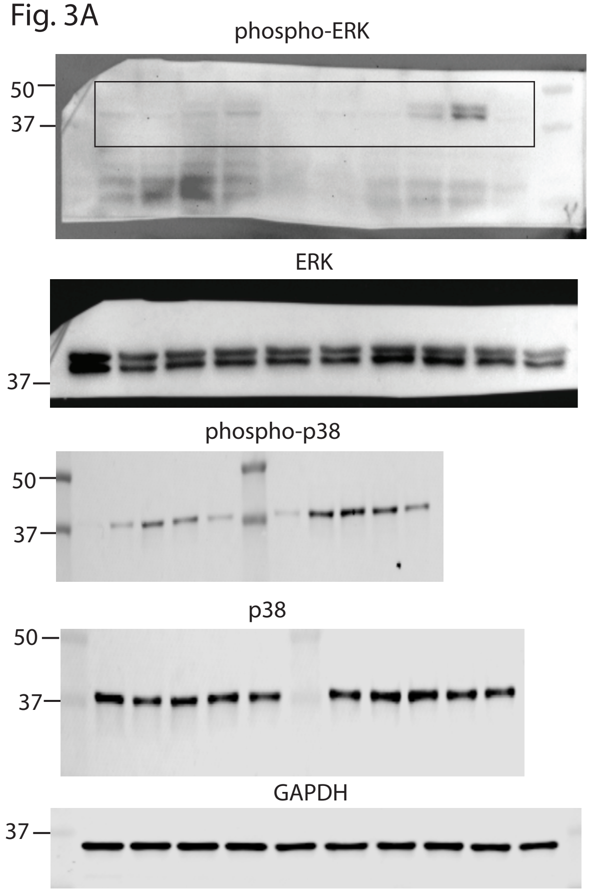


##### Fig. 3B

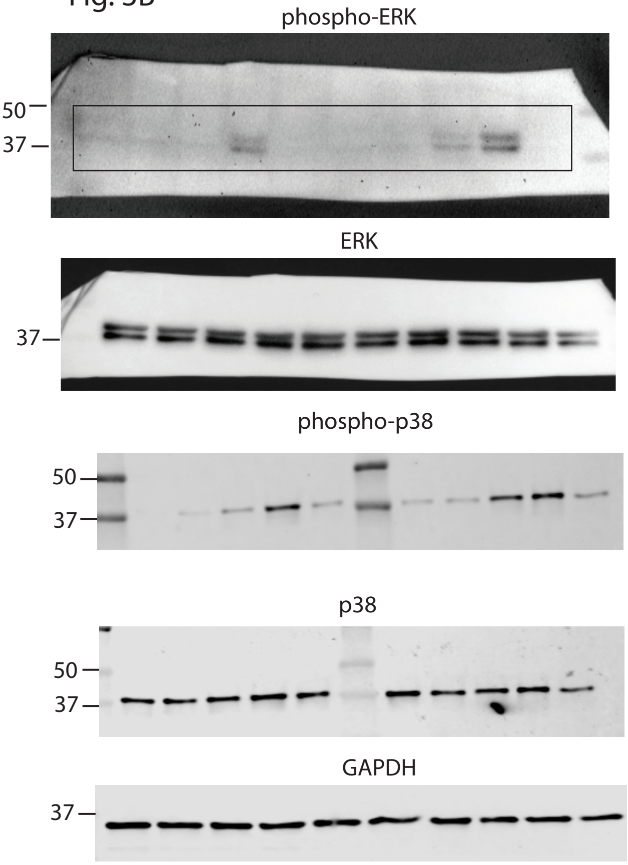

Fig. 3C

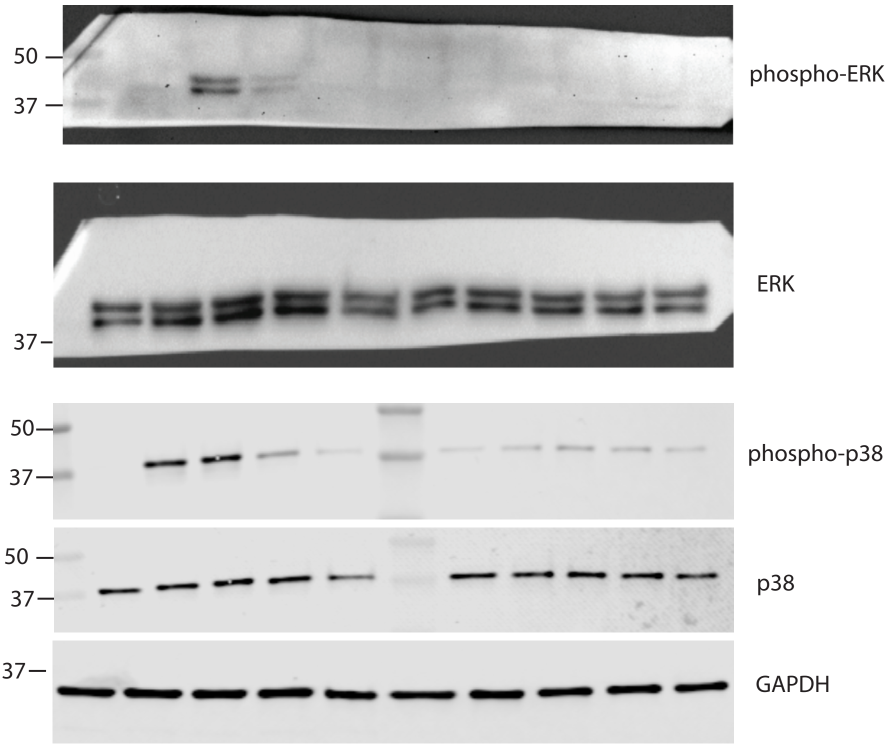


Figure 5

Fig. 5D

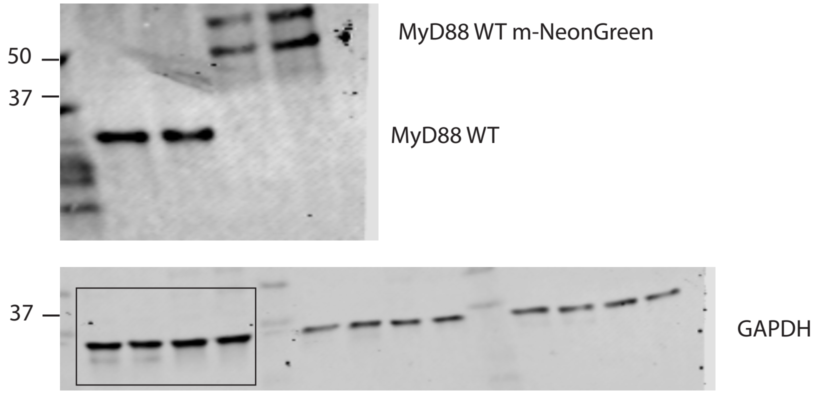
